# Integrative multi-omics identifies high risk Multiple Myeloma subgroup associated with significant DNA loss and dysregulated DNA repair and cell cycle pathways

**DOI:** 10.1101/2021.09.03.458836

**Authors:** María Ortiz-Estévez, Mehmet Samur, Fadi Towfic, Erin Flynt, Nicholas Stong, In Sock Jang, Kai Wang, Paresh Vyas, Nikhil Munshi, Herve Avet-Loiseau, Matthew W. B. Trotter, Gareth J. Morgan, Brian A. Walker, Anjan Thakurta

**Author notes:** To whom correspondence should be addressed: Anjan Thakurta, BMS, 181 Passaic Ave, Summit, NJ 07901. These authors contributed equally to this work and are co-lead authors (María Ortiz-Estévez, Mehmet Samur, Fadi Towfic).

## Abstract

Despite significant therapeutic advances in improving lives of Multiple Myeloma (MM) patients, it remains mostly incurable, with patients ultimately becoming refractory to therapies. MM is a genetically heterogeneous disease and therapeutic resistance is driven by a complex interplay of disease pathobiology and mechanisms of drug resistance. We applied a multi-omics strategy using tumor-derived gene expression, single nucleotide variant, copy number variant, and structural variant profiles to investigate molecular subgroups in 514 newly diagnosed MM (NDMM) samples and identified 12 molecularly defined MM subgroups (MDMS1-12) with distinct genomic and transcriptomic features.

Our integrative approach let us identify ndMM subgroups with transversal profiles to previously described ones, based on single data types, which shows the impact of this approach for disease stratification. One key novel subgroup is our MDMS8, associated with poor clinical outcome [median overall survival, 38 months (global log-rank pval<1×10^−6^)], which uniquely presents a broad genomic loss (>9% of entire genome, t.test pval<1e-5) driving dysregulation of various transcriptional programs affecting DNA repair and cell cycle/mitotic processes. This subgroup was validated on multiple independent datasets, and a master regulator analyses identified transcription factors controlling MDMS8 transcriptomic profile, including CKS1B and PRKDC among others, which are regulators of the DNA repair and cell cycle pathways.

**Statement of Significance:** Using multi-omics unsupervised clustering we discovered a new high-risk multiple myeloma patient segment. We linked its diverse genetic markers (previously known, and new including genomic loss) to transcriptional dysregulation (cell cycle, DNA repair and DNA damage) and identified master regulators that control these key biological pathways.

## Introduction

Multiple Myeloma (MM) patients have complex genetic heterogeneity in the tumor that includes structural variants (SVs) such as immunoglobulin heavy chain (*IgH*) translocations, single nucleotide variants (SNVs) in oncogenes and tumor suppressor genes, and genomic/chromosomal copy number variants (CNVs), as well as transcriptomic changes (1, 2). A comprehensive molecular classification of the disease based on all these types of data may shed light into how the combinations of these genetic and transcriptomic features define or contribute to intra-tumoral heterogeneity, therapeutic response and/or resistance and eventual relapse.

The MM community has devoted significant effort toward identifying molecular genetic features to diagnose MM patients, especially focused on patients with poor prognosis. For this reason, they have relied upon supervised analyses to identify molecular features associated with poor clinical outcome that may not necessarily identify biological sub-types of disease, nor be the features driving aggressive biology of the tumor. Various signatures have been previously proposed to identify high-risk patients, including UAMS70/80/17 (3), EMC92 (4), IFM15 (5), chromosome instability signature (6), centrosome index signature (7) and proliferation index (8). Some of these signatures were combined with disease stages (9) or expression of long intergenic non-coding RNAs (10) to improve their prognostic utility. Recently, we identified high-risk disease subgroups based on DNA features combining amp1q (CNV=4 or more) plus International Staging System 3 (ISS) or biallelic inactivation of *TP53* (deletion and mutation) (11); and clonal status of del17p (high-risk del17p) (12). To date, some genomic biomarkers including del17p, gain1q, t(4;14) or t(14;16), and mutations in *TP53*, in combination with clinical characteristics have been used in the clinic or clinical trials for prognosis (13, 14).

Previous efforts to stratify MM based on gene expression (GE) data identified 7 molecular subgroups with distinct transcriptomic profiles (15–17). Some of these subgroups were linked to genomic abnormalities (including translocations (SVs) or hyperdiploidy (HY)), while others such as the proliferative group (PR) apparently was driven mainly by transcriptional pathways (15). More recently, Laganà et al identified gene modules, which were subsequently associated with genomic and clinical features (17). Mutational signatures that are independent of previously defined prognostic markers have also been used to stratify MM patients (18) and stratification of MM patients based on CNVs has demonstrated some association with outcome (19).

Integrative clustering analyses across multiple data types from large, well annotated datasets, have identified novel biological subgroups in solid tumors and acute myeloid leukemia (20–23); showing the impact of data integration in disease stratification. Such an analysis, however, is yet to be reported in MM. As part of the Myeloma Genome Project (MGP) (19), here we present a large-scale multi-omics analysis of newly diagnosed MM (NDMM).

Our work identified 12 disease subgroups using an integrative multi-omics approach combining GE, SV, CNV, and SNV features (Figure 1A), where clinical covariates, such as outcome data, were not included to define genomic subgroups independently from known clinical features. We further explored the molecular features and clinical associations of the 12 biological subsets and focused on a subgroup (MDMS8) which showed the worst prognosis across the entire patient cohort (Figure 1B). MDMS8 main characteristic is a significant (>8%) genomic loss associated with dysregulated DNA repair and cell cycle/mitotic related transcriptional programs. The integrative nature of MDMS8 comes up on its transversal profile to specific known biomarkers of high risk (including 1q amplification, del17p and t(4;14) (Figure 2 and Supplementary Figure S4A-E), and to patient subgroups previously defined based only on gene expression (such as the proliferative, the MMSET and the MAF subgroups (15–17)) (Figure 6). Master regulator analysis (24, 25) identified 7 genes controlling MDMS8 transcriptional program, including E2F2, CKS1B and PRKDC, which seem to control dysregulation of DNA repair and cell cycle pathways putatively for sustaining the genome loss. We further validated MDMS8 in independent NDMM and relapsed/refractory MM (RRMM) datasets demonstrating the reproducible persistence and prevalence of this segment across patient cohorts.

**Figure 1:**
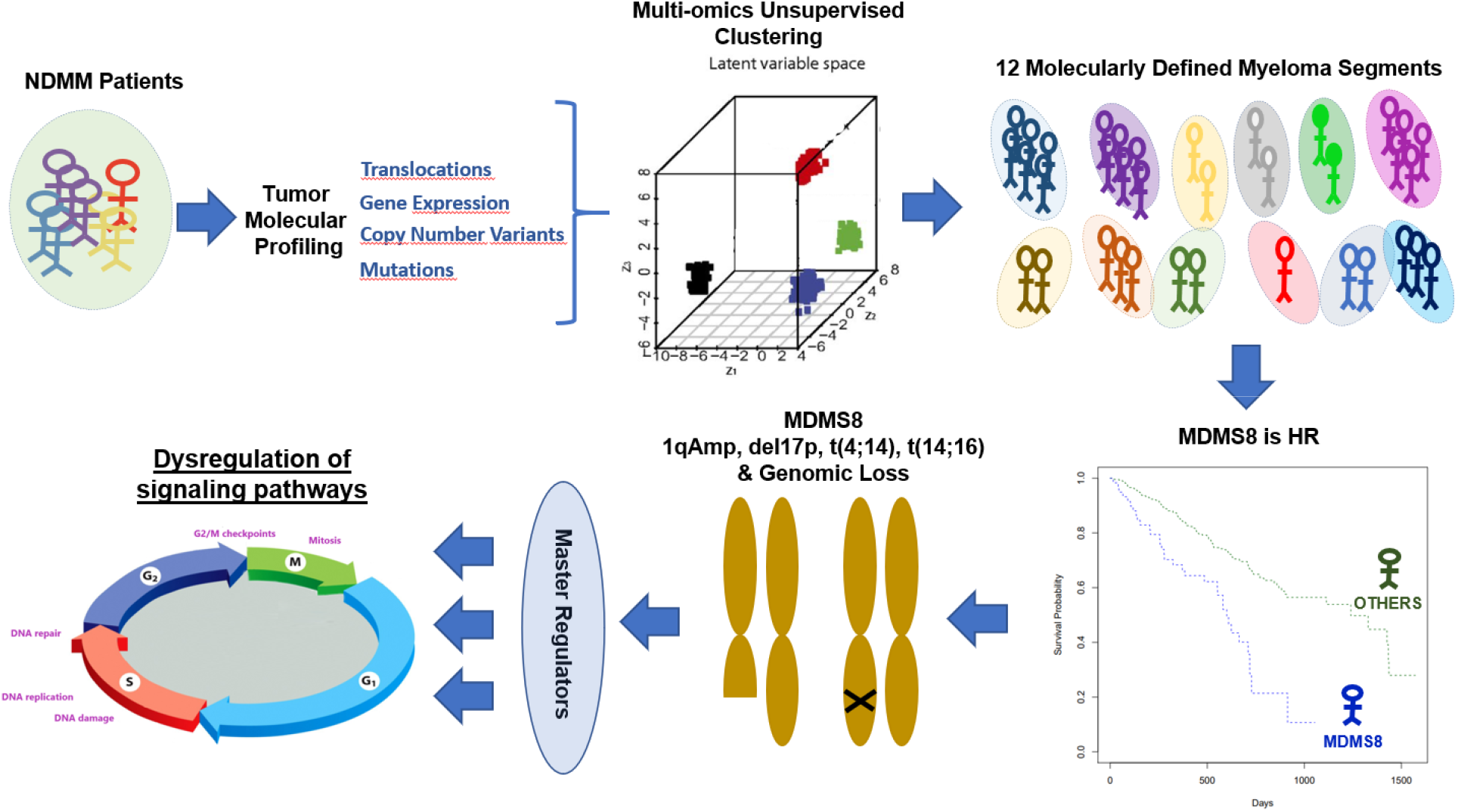
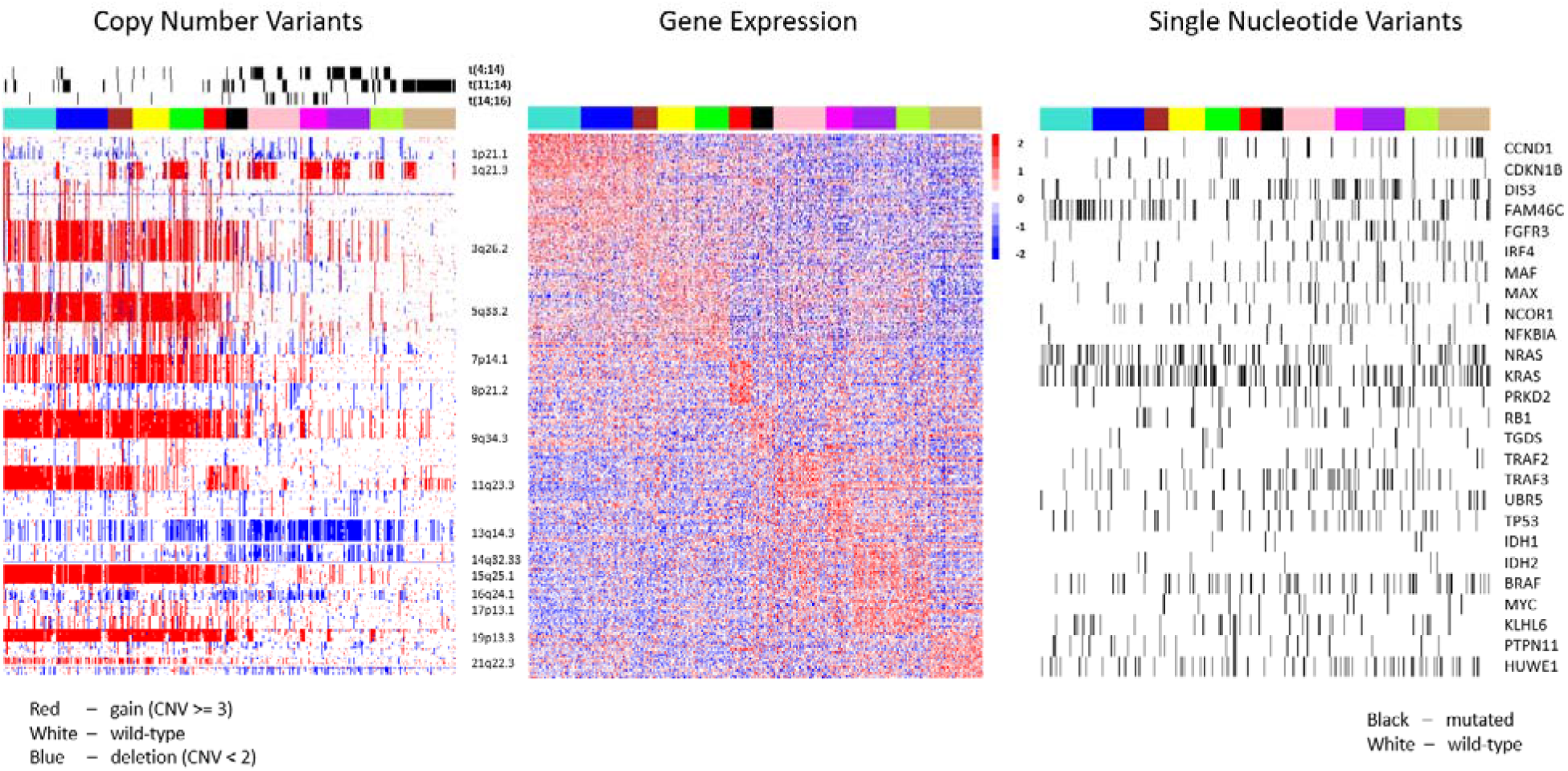
Twelve multiple myeloma subgroups identified by integrative clustering. **A)** Figure representing a visual summary of the work presented in the paper. From NDMM molecular profiles to identification of HR patient segment by multi-omics unsupervised clustering and its main characteristics including genomic loss, master regulators and DNA repair and cell cycle dysregulation. **B)** Heatmap showing molecular characteristics of the molecularly defined myeloma subgroups (MDMS 1-12): Left panel shows copy number variants with structural variants added as tracks above; middle panel shows gene expression (top 30 over-expressed genes per MDMS without replication); and right panel shows single nucleotide variants (black band denotes mutation, white band denotes wild-type sequence).

**Figure 2:**
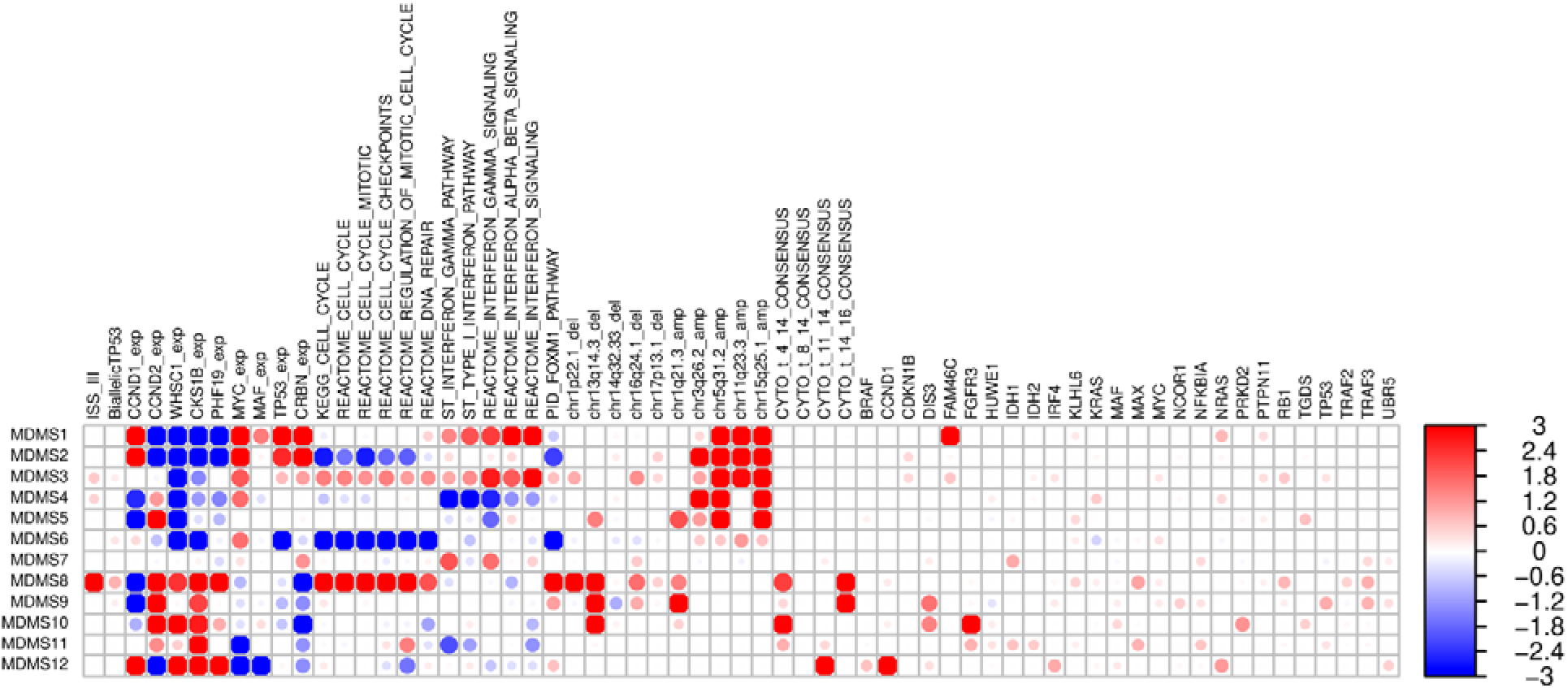
Significant genomic, transcriptomic and clinical characteristics across disease subgroups. Enrichment scores [−log10 (fdr), Fisher exact t test (binary values) or t test (continuous values) p-values]. Red and blue colors represent positive and negative associations, respectively. Values were trimmed between (−3, 3). Dot size corresponds with level of significance.

## Results

### Integrative Clustering Analysis Identifies Twelve Molecularly Defined Disease Subgroups in Myeloma

We analyzed genomic and transcriptomic data from 514 NDMM patients enrolled in the Multiple Myeloma Research Foundation (MMRF) CoMMpass study (NCT0145429, version IA17). The subset of the samples selected was based on the intersection of the various datasets (GE, CNVs, SNVs, SVs and clinical information), and patient characteristics are presented in Supplementary Table S1. Demographics, clinical data, treatment information and data processing steps have been published previously (11, 19).

Two alternative multi-omics integrative analysis methods were applied to the complete dataset: iCluster+ (26) and Cluster of Clusters Analysis (COCA) (27). Each clustering method was run one thousand times with re-sampling of features and samples to ensure robustness (Supplementary Figure S1). While iCluster+ defines clusters based on integrated, simultaneous analysis across the data types; COCA uses a two-step analysis, first clustering on each single data type and then grouping the results into a final set of clusters. Results of the two clustering methods overlapped but were not identical (Supplementary Table S2). In our dataset, iCluster+ identified 12 subgroups (in >40% of the iterations, followed by 11 clusters selected <30%) compared to 14 subgroups (>30% of the iterations, followed by 12 clusters selected <20%) identified by COCA. Consensus across iterations, defined by prevalence of same samples being clustered together, was higher in iCluster+ (>70% iCluster+ vs <65% COCA) thus, the iCluster+ output was selected for further analysis.

Twelve molecularly defined MM subgroups (MDMS) were identified by iCluster+ (Supplementary File 1), with sizes ranging from 5% to 12% of the total cohort of 514 (Figure 1B and Supplementary Figure S2). These included six HY subgroups (MDMS1-6), characterized by gains (CNV=3 or more) of chromosomes 3, 5, 9, 15 and 19, and six non-HY subgroups (MDMS7-12) (Figures 1B and 2; Supplementary Table S3). Within the HY group, MDMS1-2-3 share several molecular characteristics, including gain of Chr11 (gain11) and over-expression of *PAPD7*. MDMS1 is differentiated from MDMS3 and MDMS5 by deletion of 8p22.1 (del8p22.1), mutation of *RB1*, over-expression of *NSDHL* and up-regulated cell cycle and checkpoints signaling pathways. MDMS2 shows a deep down-regulation of cell cycle related pathways, and this characteristic is shared with MDMS6. MDMS3 is enriched in *FAM46C* and *NRAS* mutation and up-regulation of the interferon pathway. MDMS4 and MDMS5 have no gain of Chr11, but MDMS5 only is enriched in gain of Chr3 and has significant del13q and mutations in *ARID2*, *EGR1* and *NF1* genes. MDMS6 is defined by gain20q11, gain11q23.3, down-regulation of *MED11*, and down-regulation of DNA repair, cell cycle and checkpoints pathways (Figures 1B and 2; Supplementary Table S3).

Among the non-HY subgroups, MDMS7, MDMS11 and MDMS12 are significantly associated with t(11;14) (Figures 1B and 2; Supplementary Table S3). MDMS7 is also enriched in gain19q13 and up-regulated interferon pathways. Both MDMS8 and MDMS9 have t(14;16) and t(4;14) patients, however, due to the low prevalence of t(14;16) patients in the study it does not appear to be the driver of any of these groups (Supplementary Figure S3). MDMS8 is also significantly enriched in gain1q; del1p, del16q, del17p. In addition to t(14;16), MDMS9 shows a significant enrichment of gain1q, del13q14.3, del16q24.1, and mutations in *ATM*, *DIS3*, *TP53* and *TRAF3.* MDMS10 is defined by del13q14.3 and mutations in *DIS3* and *PRKD2*; while also presenting the highest significant enrichment for t(4;14) and *FGFR3* mutations compared to the other disease subgroups. The pattern of mutations in MDMS10 aligns with the activation of MEK/ERK signaling pathway (28). MDMS11 presents down-regulation of interferon related pathways (in contrast to MDMS7) and reduced expression of *FBXW2* and *KIF4B*. MDMS12, mainly driven by t(11;14), is also enriched in *CCND1*, *IRF4* and *NRAS* mutations, over-expression of *CCND1* and low expression of *CCND2* (Figures 1B and 2; Supplementary Table S3).

### Identification and Validation of MDMS8

Survival analyses were performed to understand how the molecular disease subgroups relate to clinical outcome. Eleven of the disease subgroups share a progression-free survival (PFS) and overall survival (OS) similar to standard risk patients (Figure 3) (29). In contrast, patients in MDMS8 display significantly poorer outcomes (median PFS, 19 months, log-rank p<0.001; median OS, 38 months, log-rank p<1×10^−6^) (Figure 3). MDMS8 has enrichment for ISS III patients (Fisher exact test p <0.05) and biallelic *TP53* (Fisher exact test p <0.05) (Figure 2, Supplementary Table S3). Moreover, among patients in MDMS8 carrying previously described high-risk markers in MM, including t(4;14), t(14;16), gain1q, del13q and del17p, both PFS and OS are significantly worse than among patients with similar genomic characteristics in non-MDMS8 clusters (Figure 4). Separate analyses for each of these high-risk markers, showed similar results, suggesting the presence of a common biology across these different genomic groups in addition to their high-risk features contribute to overall clinical outcome (Supplementary Figure S4A-E).

**Figure 3:**
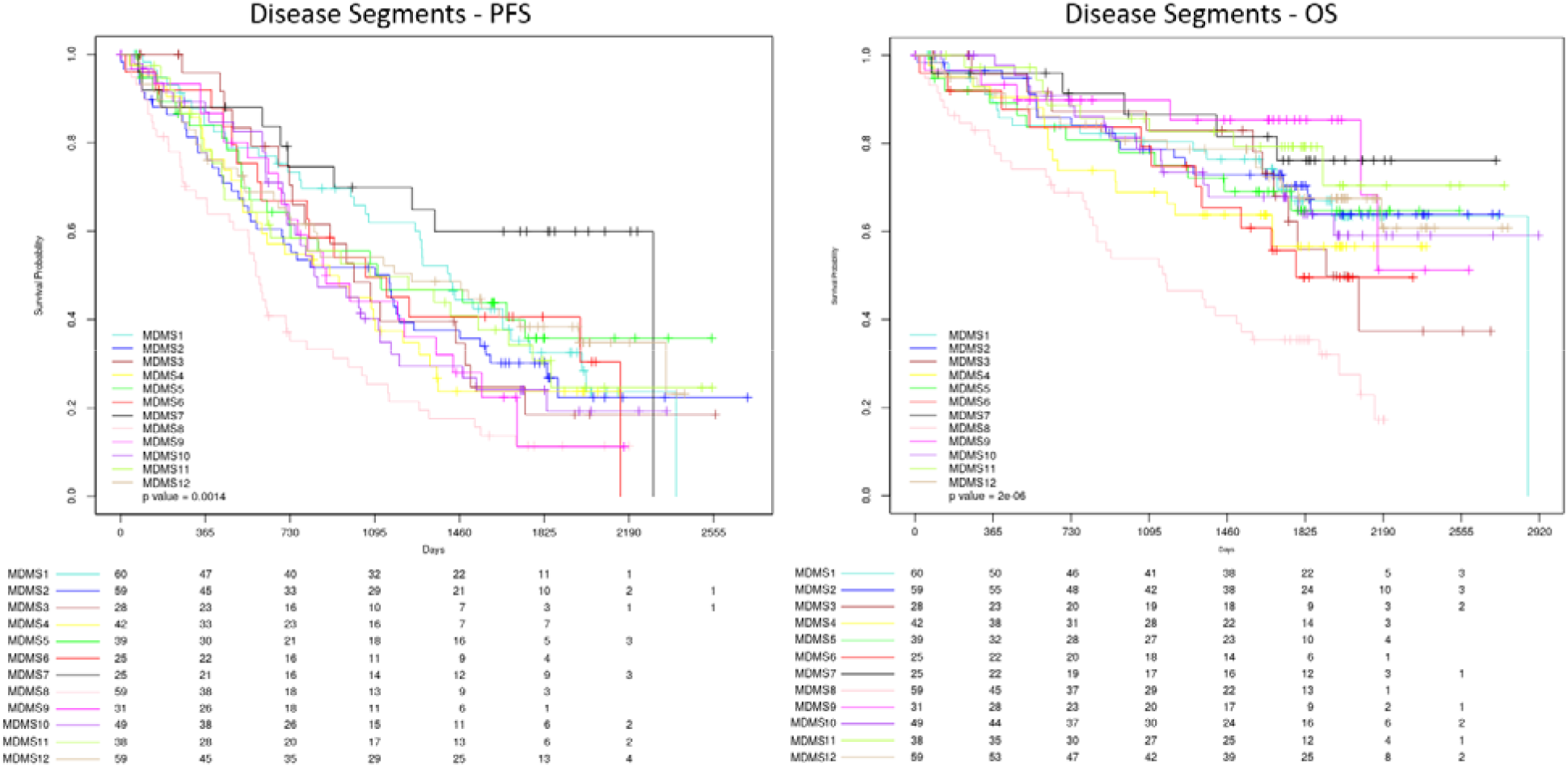
Kaplan-Meier (KM) survival analysis of outcome among the disease subgroups MDMS 1-12. Progression-free survival (left) and overall survival (right) among patients in each of the 12 myeloma subgroups. Global log-rank p-value shown for each KM plot.

**Figure 4:**
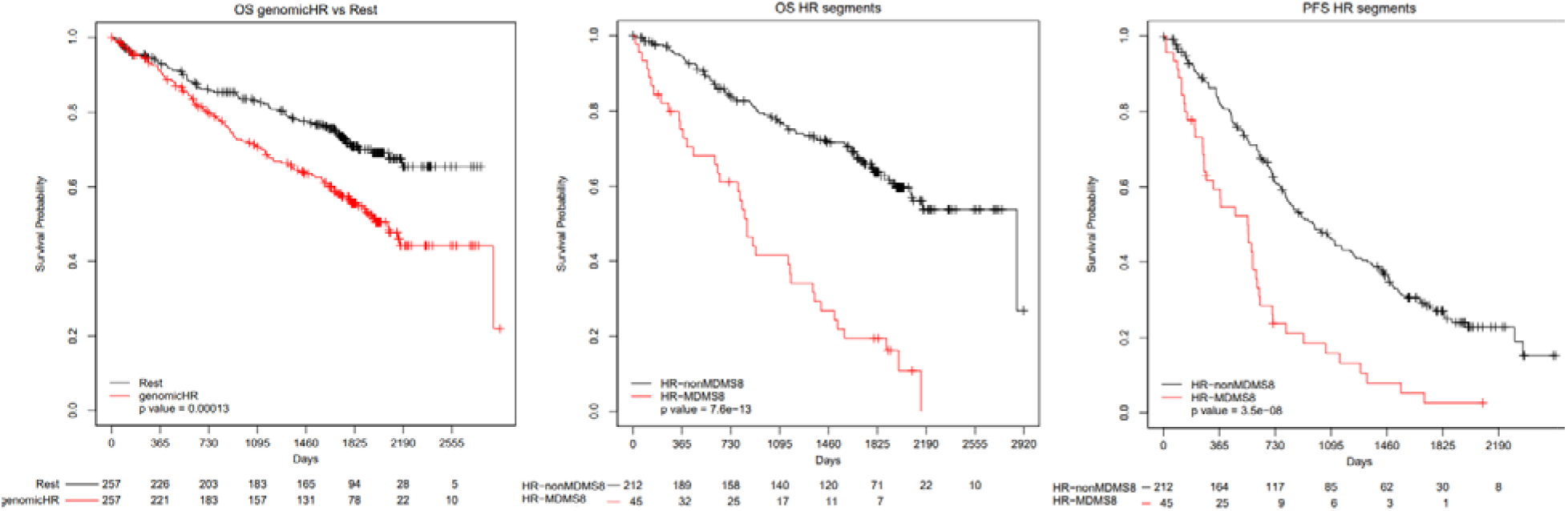
Kaplan-Meier (KM) survival analysis of genomic subgroups versus MDMS8. KM survival analysis showing overall survival (OS) of patients carrying one or more of the following genomic aberrations: [t(4;14), t(14;16), gain1q or del17p] versus the remaining patients (left); overall survival (OS) of patients with genomic aberrations [t(4;14), t(14;16), gain1q or del17p] in MDMS8 versus the same subset of patients in non-MDMS8 subgroups (middle); and progression free survival (PFS) of patients with genomic aberrations [(t(4;14), t(14;16), gain1q or del17p] in MDMS8 versus the same subset of patients in non-MDMS8 subgroups.

In MDMS8 patients, DNA repair/damage related genes, such as *ARID2*, apoptosis related *BIRC2*, *TRAF1*, *TRAF2* (30, 31), and genes associated with CDK function, including *MAX*, *RB1*, and *TP53* (32, 33), are significantly mutated. Differential GE analysis identified significant activation of genes controlling mitotic and DNA damage/repair processes (*CENPI*, *SKA1*, *NUF2*, *PLK1*, *AURKB*, *BIRC5* and *BUB1*), DNA synthesis (*POLA1*, *PRIM1* and *PRIM2*), and checkpoints (*MCM/CDC/RFC* gene families and *CDK1/2*)-all generally involved in cell cycle related pathways (Figure 5A). A differential gene expression analysis comparing patients with shared genomic characteristics (including t(4;14) or gain1q) in MDMS8 versus non-MDMS8 patients shows DNA repair, mitotic, checkpoint and *MYC* pathways significantly up regulated in MDMS8 (Supplementary Figures S4A-B).

**Figure 5:**
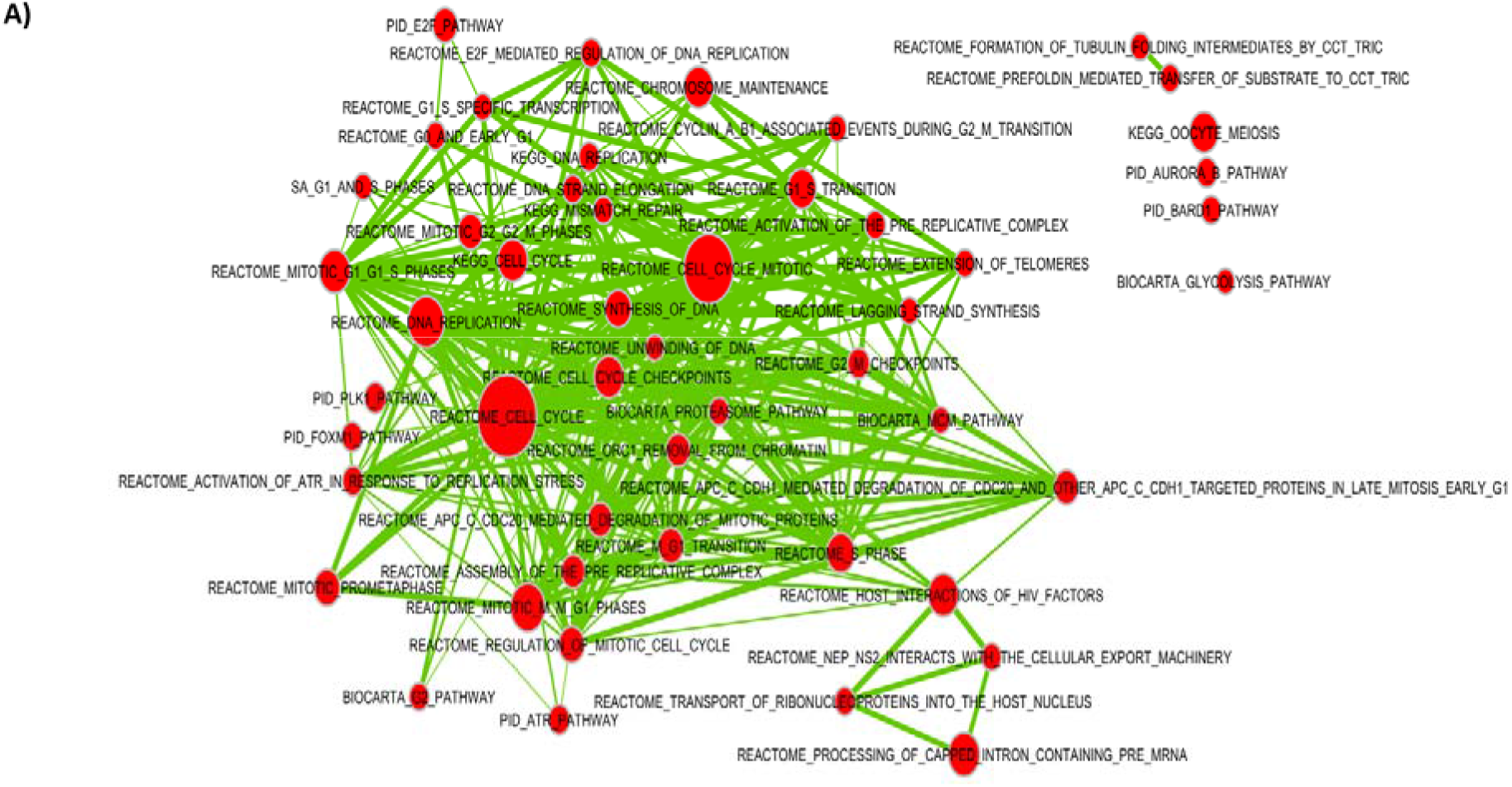
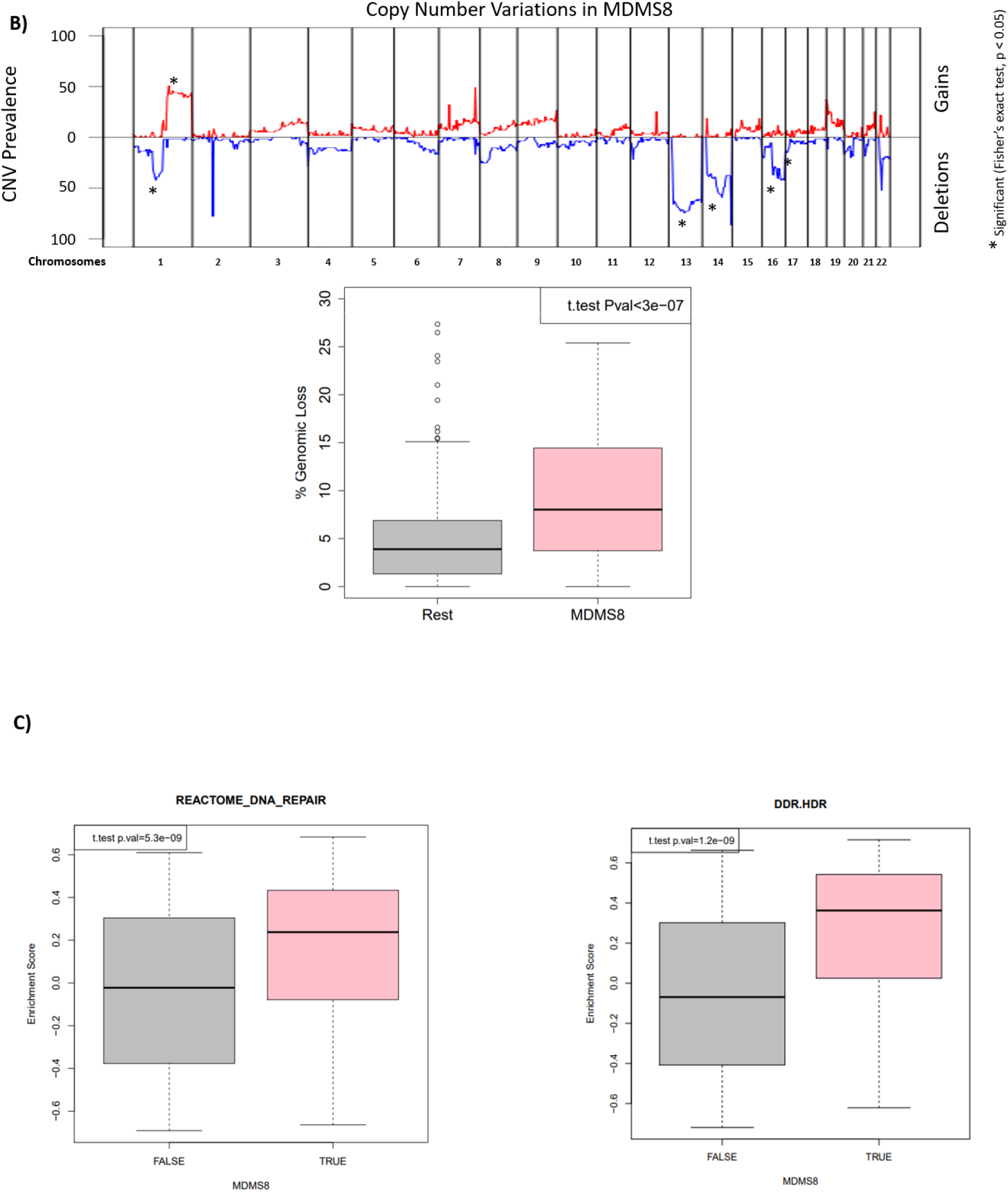
Genomic and gene expression characteristics of MDMS8. **A)** Signaling pathway network showing significantly up-regulated pathways in MDMS8 compared to the rest of the disease subgroups. **B)** Prevalence of deletions (negative Y-axis, blue) and gains (positive Y-axis, red) across the genome in MDMS8 (top panel). Percentage of genomic losses in MDMS8 vs the rest of ndMM patients (bottom panel). **C)** Enrichment scores of the Reactome DNA repair pathway in MDMS8 vs the rest (left panel) and Homology-dependent Recombination (HDR) pathway in MDMS8 vs the rest (right panel).

The genomes of MDMS8 samples present an increased loss of genes on various chromosomes, including 1, 13, 14, 16 and 17 on the p arm (Figure 1 and top panel of Figure 5B) compared to the other molecular subgroups. We calculated the number of genomic cytobands containing a deletion and the total amount of genomic deletion in all samples (measured by the extent of deletion as percentage of the whole genome), which showed a significantly increased number of genomic regions having a loss in MDMS8 (median > 8% of genomic loss (Methods)) compared to the rest of the patients (median < 4% of genomic loss) (t.test p.val < 1e-6, bottom panel Figure 5B). A gene set variant analysis (GSVA, see methods) on DNA damage/repair pathways (including REACTOME and DNA Damage Response (DDR) pathways (55)) showed a significant up-regulation of REACTOME DNA damage and repair pathways, as well as the DDR Homology-dependent recombination (HDR), Translesion Synthesis (TLS) and Base Excision Repair pathways in MDMS8 compared to the other NDMM patients (Figure 5C, Supplementary Figure S5).

To explore the prevalence of MDMS8 in other MM datasets, we built a GE classifier on the discovery data, applied it to independent cohorts (including IFM (5) and APEX (15, 35) (Supplementary Figure 6A), and UAMS (17) (Figure 6)), and explored prevalence and genomic properties (when available) of patients classified as ‘MDMS8-like’ (Supplementary Figure 6B). We generated a multiclass linear model classifier with lasso regression for feature selection based on gene expression, since it was the common datatype available across the datasets. The trained classifier comprised a linear model on the expression of 35 genes (Supplementary Table S4). The training performance of the classifier for MDMS8 has a recall ~80% and precision of 75% (where false positives were mostly patients from MDMS9 and MDMS10) (Supplementary Table S5). Information on the training performance of the classifier for all clusters is shown in Supplementary Table S5, with a median recall of 60% and precision of 64%; where most of the mis-classified calls happened between HY groups. Application of the classifier to the IFM dataset (Supplementary Table S3) identified a MDMS8-like group with similar prevalence (~12%) and significantly poorer OS (median OS not reached, long rank p < 1e-4) (left panel of Supplementary Figure S6A). Importantly, the MDMS8-like group in IFM also presented the high rate of genomic loss (median genomic loss MDMS8-like >8% and rest < 4%, Supplementary Figure S6B), validating not only the gene expression profile but also the genomic features. We applied the classifier to the APEX trial Affymetrix-based GEP dataset (RRMM) (15, 35), where, again, there was a significant difference in OS observed between MDMS8-like versus other RRMM patients (right panel of Supplementary Figure S6A). Prevalence of the MDMS8-like segment in the APEX trial was <15%. This analysis demonstrates that MDMS8-like segment is reproducible across multiple datasets and that its poor OS is independent of treatment regimen.

**Figure 6:**
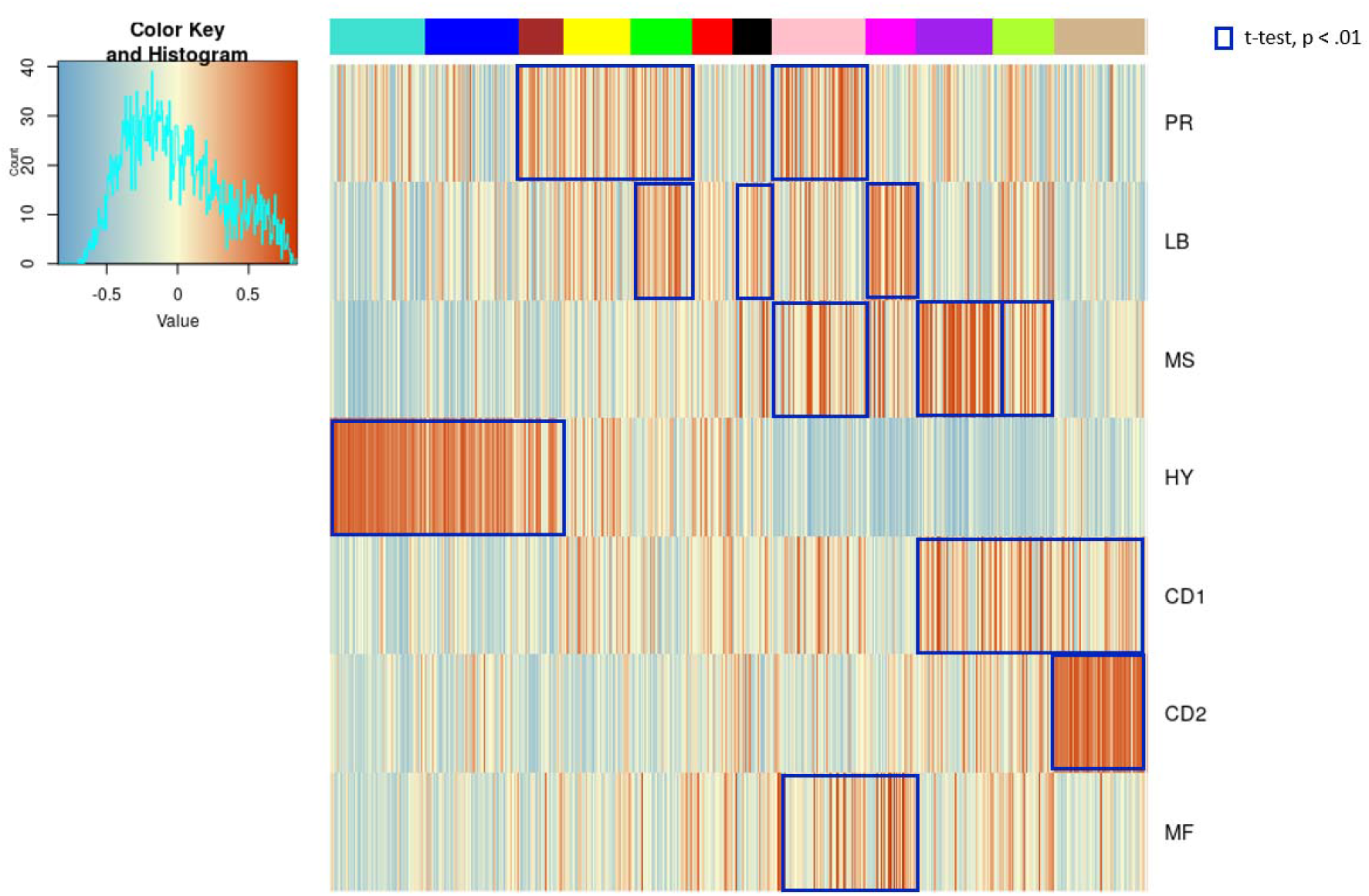
Comparison of MDMS8 to other Gene Expression Signatures. **A)** Gene expression enrichment of Zhan et al GE patient subgroups signatures (15) across the twelve molecularly defined myeloma subgroups. Red represents positive enrichment; blue represents negative enrichment. Blue squares highlight significant association (enrichment scores t-test p<0.01) between the Zhan et al signatures and MDMS disease subgroups.

### MDMS8 Comparison to Previously Reported MM Subgroups and High-risk Signatures and Biomarkers

To place our analysis in the context of previous efforts, we explored similarities and differences between MDMS8 and other MM subgroups identified using GE datasets by Zhan et al (15) and Broyl et al (16). In Figure 6 (and Supplementary Figure S7) MDMS8 shows a significant enrichment in the signature scores of the publicly described PR (proliferative), MS (MMSET) and MF (MAF) groups, which is coherent with MDMS8 since it contains t(4;14) patients (MS group), t(14;16) patients (MF group) and it shows dysregulation of cell cycle (PR group). Conversely, Zhan et al groups are associated with multiple MDMS clusters, suggesting no 1:1 association between the two clustering approaches. We also applied our classifier to the Zhan et al GEP discovery dataset and compared our cluster calls to theirs. This comparison, again, shows commonalities among some of the groups, such as the HY (hiperdiploid) from Zhan et al which contains most of our MDMS3 and MDMS5, while CD2 maps uniquely to MDMS12; but it also shows clear differences, including MDMS4 (which from our genomics data is HY) which doesn’t associate to the previously defined HY group. Also, MF and MS groups are subdivided into various MDMSs. Finally, MDMS8, presents a transversal profile to the previously defined GEP subgroups (containing patients from MF, MS, MY and PR) suggesting the biology of this group is more heterogeneous than what was previously described (Supplementary Table S6). While both attempts (Zhan et al and ours) are unsupervised in nature, results show key differences between using GE only vs multi-omics integrative approach. Comparison of MDMS8 with the CNV clusters defined by Walker et al (19) identifies significant enrichment of CN7 (characterized by gain1q and del13q); however, the CN7 cluster does not include all of the MDMS8 patients, notably excluding those with t(4;14).

UAMS70 (3) and EMC92 (4) high-risk MM classifiers were applied to the discovery dataset to explore the overlap between patients deemed high-risk by these outcome-based classifiers and MDMS8 patients. MDMS8 captures a significant number of high-risk patients identified by both EMC92 (34%) and UAMS70 (40%). A third of MDMS8 patients, however, were not captured by these high-risk GE-classifiers (Supplementary Figure S8). Discordance among these groups is not unexpected, given that the number of shared genes between UAMS70 and EMC92 signatures is <5%. Moreover, unlike the GE-classifiers, the unsupervised approach used to identify MDMS8 was not based on clinical outcome.

### Master Regulators Drive Transcriptional Phenotype in MDMS8

Finally, a master regulator (MR) analysis using msVIPER (36) was performed to elucidate the mechanisms linking genomic alterations to the transcriptional profiles of MDMS8. The master regulator genes were selected on the basis of impact on transcriptional changes of their inferred downstream targets (regulons) using a context-specific gene regulatory model (37). Ten MRs were identified (Figure 7 and Supplementary Figure S9), with seven of them showing positive activation in MDMS8: *E2F2*, a transcription factor member of the e2f family; *CKS1B*, a protein kinase regulator located in 1q21; *RBL1*, which encodes a gene that is similar in sequence and possibly function to retinoblastoma 1 (*RB1*), significantly mutated in MDMS8; *PRKDC,* a protein kinase sensor for DNA damage incurred in DNA repair/recombination; *RUSC1,* related to the Trk receptor signaling mediated by the MAPK pathway; *NUP93*, described as tumor growth modulator via cell proliferation and actin cytoskeleton remodeling (38) and migration and invasion capacity of cancer cells (39), and *MSN,* Moesin, described as an unfavorable prognostic biomarker in various cancers (40–42). Genes encoding the two zinc finger proteins (*ZBTB40* and *ZNF837*) and the histone deacetylase 3 (*HDAC3*) were down-regulated MRs (Supplementary Figure S9).

**Figure 7.**
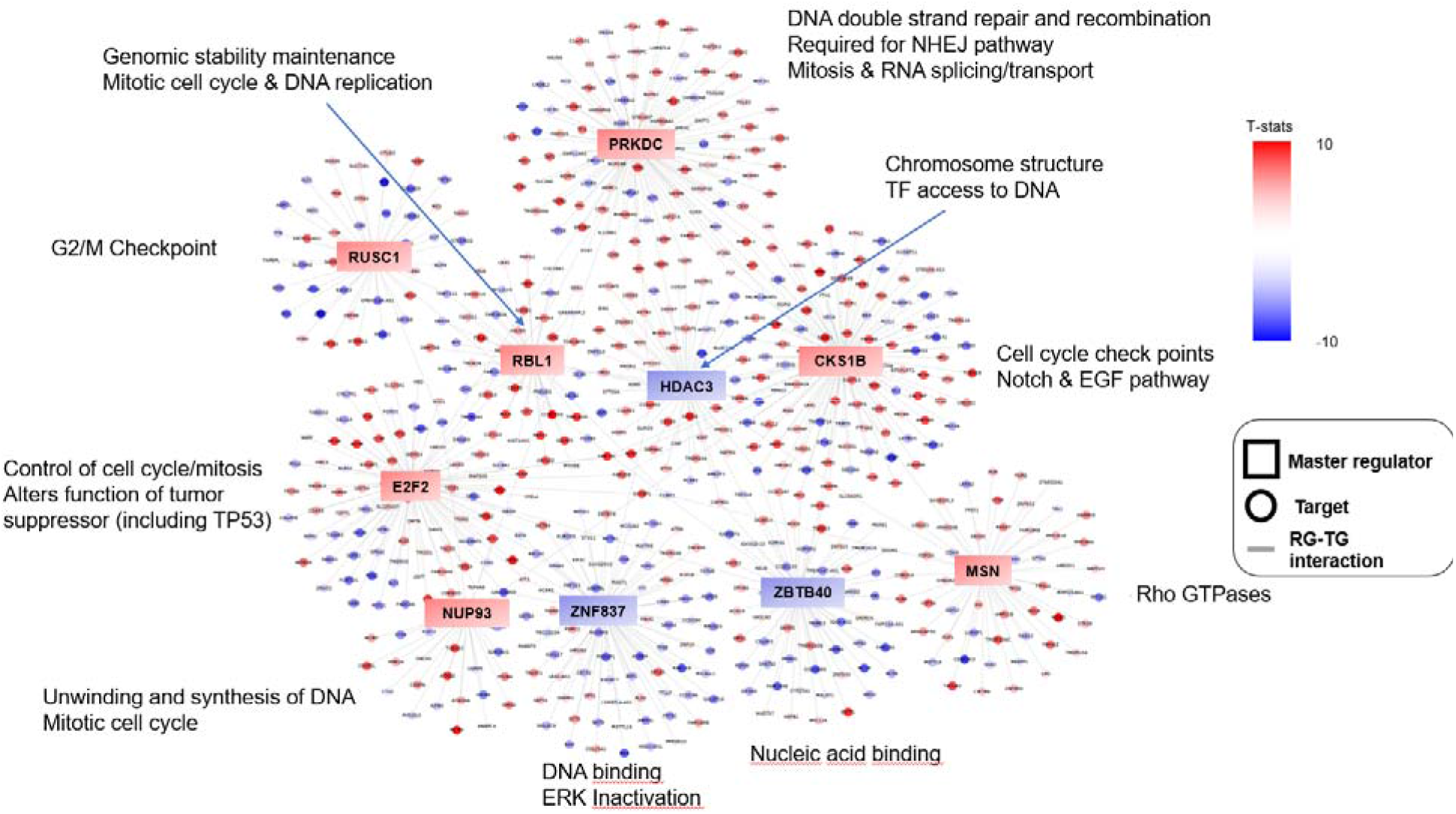
Master regulator analysis. Master regulators’ regulons and their associated signaling pathways. Color scheme represents −log10 (t test p-value) of activation score of the listed genes in MDMS8 versus the rest, with red for positive values and blue for negative values. Squares represent master regulators; circles represent regulon genes.

An enrichment analysis based on the regulons of MDMS8 MR was performed to understand MDMS8 biology and signaling functions controlled by these MRs. Most of the activated MRs control diverse biological processes (Figure 7) including ones related to mitosis, such as the *E2F2* regulon, which contains the *KIF* family, and the *CKS1B* regulon with *RAD21* and the *MCM* family; or the *MSN* regulon, associated with Rho GTPases (switches that regulate the actin cytoskeleton, influence cell polarity, microtubule dynamics, membrane transport pathways and transcription factor activity (43)). Cell cycle and DNA repair pathways in MDMS8 appear to be controlled by *RBL1*, *NUP93* and *PRKDC*, although genes in the PRKDC regulon are involved also in spliceosome and RNA transport pathways, consistent with MDMS8 biology.

Regulons downstream of the negatively activated MRs were not significantly associated with any specific signaling pathways, although they contained previously defined tumor suppressor genes, such as *KDM4A* (44) and *E2F4* (45). Of the MRs, the specific roles of PRKDC and RBL1 and their regulons in DNA damage/repair would be consistent with supporting the maintenance of MDMS8 myeloma cell’s loss of genetic material.

## Discussion

In this study, we describe molecular segmentation of NDMM by a joint modeling of multiple omics data types to identify common latent variables to group patient samples into biologically distinct disease subtypes. Our unsupervised analysis identifies twelve biological subgroups of MM, confirming hyperdiploidy-dependent and SV-dependent as the two predominant molecular subtypes of MM. Notably, we identified and replicated a new disease segment (MDMS8) that is enriched in diverse known high-risk genomic features, accompanied by various MM driver mutations and dysregulation of DNA damage and repair pathways and cell cycle/mitotic processes, alongside a genome loss, that had not been previously described in MM. Master regulator analyses identified potential drivers of the transcriptional program pointing to key pathways in DNA repair, cell proliferation, cell cycle progression and chromosomal stability and maintenance. PFS and OS are significantly inferior for patients in MDMS8 compared with patients in non-MDMS8 subgroups, even when patients in both cohorts carry the same high-risk genomic biomarkers, including 1q gain, del17p, t(4;14) and/or t(14;16). Our analysis shows for the first time that along with the different high risk markers (del17p, t(4;14), amp1q) in ndMM there is a common transcriptional program linked to the accumulation of genome loss in a subset of those tumors. In our estimation, the identification of MDMS8 by the integration of multiple data-types enabled a transversal and improved molecular description of high risk MM biology over previous GE-based or CN-based approaches. Not surprisingly, due to its association with poor clinical outcome, MDMS8 contains a significant number of patients picked up by gene expression based high-risk classifiers, EMC92 (4) and UAMS70 (3). Besides, our integrated clustering analyses separate t(4;14) MM samples into multiple disease subgroups, including MDMS10 and MDMS8, all with high MMSET/NSD2 expression independent of the disease segment. The outcome and transcriptomic profile of MDMS8, however, are distinctly different from patients with t(4;14) in other disease subgroups, suggesting that overexpression of MMSET/NSD2 per se does not play a direct role in high-risk biology as had been previously discussed in the literature. While additional work is needed to tease out the implications of such observations, taken together, our results suggest that an integrated analysis of multiple data types could effectively sort out the heterogeneity of t(4;14) myeloma.

Identification of MDMS8, and its genomic loss linked with the dysregulated transcriptional phenotype prompted our exploration of functional drivers. The mechanism of the genome loss or its association with high-risk genetic loci is not clear at this time. Gene set enrichment analysis however revealed the relationship between MDMS8 transcription profiles with DNA repair/damage and cell cycle pathways, especially those directing the mitotic machinery and steps required for functional cell division. We envision that MDMS8 cells have adaptive mechanisms to tolerate excess DNA damage. It is likely that these transcriptional pathways are critical for repairing DNA damage as a consequence of DNA replication or induced to relieve the stress of multiple steps of proper chromosomal segregation during mitosis. All 7 MRs whose activities are up-regulated in MDMS8 are essential genes in MM, controlling key biological functions required for DNA repair/damage, cell cycle check points for G1/S and G2/M, MYC-driven growth and survival pathways and mitotic processes. This analysis provides a pool of proteins to potentially target the underlying biological basis of the aggressive nature of the disease. Similar approaches in other cancers (24) have suggested possible synthetic lethal relationships between MRs which could provide novel combination approaches for therapeutics development in high-risk MM. These efforts could be combined or complemented with targeting the dysregulated DNA damage repair pathways.

In conclusion, this work presents an integrative clustering-derived molecular classification of Multiple Myeloma using key genetic features with the transcriptome. We find a molecular segment enriched in extensive DNA loss, accompanied by upregulated DNA damage repair and cell cycle/mitotic pathways. This integrative analysis also illustrates that this type of approach could improve our understanding of the disease heterogeneity of Multiple Myeloma by studying the individual molecular segment such as MDMS8.

## Methods

### Data processing

#### Gene expression

RNA extraction, library preparation and sequencing for both MMRF CoMMpass and IFM/DFCI were previously described by Walker et al (19) and https://research.themmrf.org.

#### BAM to FastQ file conversion for MMRF CoMMpass cohort

Previously aligned BAM files were collected from database of Genotypes and Phenotypes (dbGaP) and converted to FASTQ using Picard tools v2.1.1 to extract read sequences and base quality scores.

#### Quantification

FASTQ files from both cohorts were quantified using Salmon. Isoform level expressions were quantified with Quasi-mapping using GRCh38 cDNA reference genome from Gencode v24. Gene level abundances were calculated using tximport and isoform level TPM (transcript per million) estimates for each sample.

#### Affymetrix gene expression

GE data coming from Affymetrix HG-U133 Plus 2 were normalized using EdgeR (46) package available in CRAN.

#### Scaling gene level expressions and selecting high variable genes

GE was normalized for each sample against three housekeeping genes. 11 housekeeping genes (47) were originally tested and the top 3 genes with lowest standard deviation were selected. Geometric mean of these 3 housekeeping genes (NONO, PGK1 and VPS29) was used to scale gene level expressions.

#### Calling copy number variants

preprocessing for copy number analysis has been described previously Walker et al (19). Genomic loss was calculated in each sample adding all the length of all the subgroups with a “loss” call from control-freec output (including both homozygous and heterozygous deletions). The final proportion of genomic loss is calculated per patient using size of genomic loss previously calculated over the genome size.

#### SNV data

SNVs were called and preprocessed as previously described (19). After preprocessing, only missense mutations that were observed in ≥ 3% of the patients were kept for further analysis.

#### SV data

SVs were called and preprocessed as previously described (19). Lowly prevalent SVs might be under-represented in our dataset due to size limitations.

### Clustering

Two different clustering algorithms iCluster+ (26) and the Cluster of Clusters Algorithm (COCA) by the Cancer Genome Atlas Research Network (27) that integrate multiple OMICs data types with different approaches were run with a range of parameters to identify the combination which produced the most robust and stable clusters across our dataset. The number of clusters ranged between 2 and 20, and the optimal solution was selected based on Bayesian Information Criteria (BIC). Membership consistency across iterations was used to select iCluster+ as the final clustering approach. More information can be found in the Supplementary Methods File.

### Biomarker Analysis

#### Differential Gene Expression

Voom-LIMMA was run for GE analysis, using linear models to assess differential expression in the context of multifactor designed experiments (49). It was implemented in the *limma* package for Bioconductor (http://www.bioconductor.org) and applied to test differential relative abundance between conditions for each cluster independently. Significance p-values were corrected for multiple testing by the false-discovery method and deemed significant at an FDR threshold of 0.05 (5%) (50).

#### Pathway Analysis

Gene-set enrichment analysis (GSEA (51)) was applied to rank relative abundance ratios obtained during differential analysis for each comparison. Weighted enrichment statistic calculations were used instead of the classic unweighted ranking to account for fold change differences in addition to protein ranking. Gene categories assessed for enrichment corresponded to the canonical pathway collection (e.g. Reactome, Biocarta, KEGG) obtained from the *MSigDB* database (file: c2.cp.v5.2.symbols (52)). Enrichment p-values were corrected for multiple testing by FDR.

#### Signature Enrichment Analysis

GSVA r package was used to calculate enrichment analysis of the various signatures. For UAMS70 (3) and EMC92 analysis, thresholds were refined to RNAseq data to select respectively 15% and 20% of the population with the highest scores.

#### Identification of master regulators

Master regulator analysis was performed using the msVIPER algorithm in the VIPER R package. More information can be found in the Supplementary Methods File.

#### Classifier

we utilized the glmnet package in CRAN (https://cran.r-project.org/web/packages/glmnet/index.html) to estimate a multinomial elastic net modelregression model with cross validation. The features were selected by estimating models with 100 nfolds on the top 3813 genes by coefficient of variance across all datasets. 42 genes identified across the cross-validation iterations were included in the final model. In order to make all the MM datasets comparable, they were normalized together with voom/limma and dataset bias was removed with Combat R function (53). Finally, all datasets were scaled independently by genes to median=0 and standard variation = 1.

#### Statistical analyses

various statistical tests from the stats v3.5.3 R (54) CRAN package were used to check significance of the association of the subgroups to different variables. Fisher’s exact test for binary data (mutations/CNVs), t-test for continuous variables (GE pathway scores), and global log-rank test for outcome (PFS/OS).

## Data Availability

Sequencing data were deposited in the European Genome Archive under accession EGA00001001147 and EGA00001000036 or at database of Genotypes and Phenotypes (dbGAP) under accession phs000748.v5.p4.

## Code Availability

Our genomic pipeline code is provided under https://github.com/celgene-research/mgp_ngs. Methods used for analysis are publicly available.

## Acknowledgements

The authors acknowledge continued support for MGP from colleagues at BMS, especially Dorothy Fallows, Rupert Vessey, Douglas Bassett, Amit Agarwal and the Myeloma Disease Strategy Team.

## Authorship Contributions

The project was conceived and designed by AT. Funding acquisition by EF and AT. Project administration by MO, FT and EF. Oversight and management of resources (data generation, collection, transfer, infrastructure, data processing) by EF, FT, MS, BW, NM, HA-L, AT, GJM. Analyses and interpretation were designed and performed by MO, FT, MT, NS, MS, EF, IJ, KW, BW, PV, HA-L, GJM, NM, AT. Data visualization performed by MO, MS, and FT. Supervision and scientific direction provided by AT. The manuscript was written by MO, MS, FT, EF, AT.

