## Supplementary for "Integrative multi-omics identifies high risk Multiple Myeloma subgroup associated with significant DNA loss and dysregulated DNA repair and cell cycle pathways"

### Supplementary Document

María Ortiz-Estévez[1†], Mehmet Samur[2†], Fadi Towfic[3†], Erin Flynt[4], Nicholas Stong[4], In Sock Jang[3], Kai Wang[3], Paresh Vyas[5], Nikhil Munshi[6], Herve Avet-Loiseau[7,8], Matthew WB Trotter[1], Gareth J. Morgan[9], Brian A. Walker[10] and Anjan Thakurta[4*]

[†]These authors contributed equally to this work and are co-lead authors.

1 BMS Center for Innovation and Translational Research Europe (CITRE), A Bristol Myers Squibb Company, Sevilla, Spain

2 Dana-Farber Cancer Institute, Harvard TH Chan School of Public Health, Boston, MA, USA

3 Bristol Myers Squibb, San Diego, CA, USA

4 Bristol Myers Squibb, Summit, NJ, USA

5 Molecular Haematology Unit, BRC Haematology Theme, Oxford Biomedical Research Centre, Oxford Centre for Haematology, Weatherall Institute of Molecular Medicine, Radcliffe Department of Medicine, University of Oxford, Oxford, UK

6 Dana-Farber Cancer Institute, Harvard Medical School, Boston, MA, USA

7 IUC-Oncopole and Cancer Research Center of Toulouse, France

8 INSERM U1037, Toulouse, France

9 New York University, New York, NY, USA

10 Melvin and Bren Simon Comprehensive Cancer Center, Indiana University, Indianapolis, IN, USA

### Table of Contents


**Supplementary Methods** **3-5**

**Supplementary References** **5-6**

**Supplementary Figure Legends** **7-8**

**Supplementary Figures**

*Figure S1*. Analytic Methodology **9**

*Figure S2*. Consensus Matrix and Joint Variation Analysis **10**

*Figure S3.* MAF expression **11**

*Figure S4.* Enrichment scores of DDR pathways in MDMS8 vs rest **12-16**

*Figure S5 A-E*. Genomic Markers in MDMS8 vs non-MDMS8 **17-18**

*Figure S6 A-B*. MDM8-like in IFM and APEX. Genomic loss of MDMS8-like in IFM **19**

*Figure S7*. Comparison of MDMS to Segments from Broyl et al **20**

*Figure S8*. Comparison of MDMS8 to HR Gene Expression Signatures **21**

*Figure S9*. Master regulators of high-risk patient segment MDMS8 **22**

**Supplementary Table Legends** **23**

**Supplementary Tables**

*Table S1*. Dataset Characteristics **24**

*Table S2*. COCA vs iCluster+ segments **25**

*Table S3*. Cohort and Genomic Features of MMDSM1–12 **26**

*Table S4*. Classifier features **26**

*Table S5.* Confusion matrix of cluster labels and classifier predictions on the discovery dataset **27**

*Table S6.* Comparison of MDMS8 to the previously defined GE UAMS groups **28**

### Supplementary Methods

***Datasets:*** The Myeloma Genome Project (MGP) is a collaborative research initiative to gather and uniformly analyze genetic datasets that have been generated on samples obtained from newly diagnosed multiple myeloma (NDMM) patients [1]. Next generation sequencing (NGS) data from patients with NDMM were processed and analyzed in a uniform manner as described by Walker et al [1]. For this analysis, patients with available baseline data, including whole exome and genome sequencing (WES and WGS), RNA sequencing (RNAseq), progression free survival (PFS), and overall survival (OS, median OS was not reached in any of the datasets) (n=514) were extracted from the full MGP dataset (N=1273). Cohort characteristics of the full MGP population and the subset used in this analysis are summarized in Supplemental Table S3.

Data from 344 newly diagnosed MM (NDMM) patients from the IFM portion of the IFM/DFCI2009 clinical trial [2], from whom we had RNAseq, were used for validation (ClinicalTrials.gov Identifier: NCT01191060). One hundred of these had the full set of data types used for the initial clustering.

The APEX trial dataset [3] (NCT00048230) includes patients with measurable progressive MM after one to three previous treatments. Patients were treated with either bortezomib or high-dose dexamethasone. This dataset was used for validation of the high-risk group in the relapse setting.

The UAMS dataset (GSE2658) includes CD-138-selected plasma cells from bone marrow of 559 patients with newly diagnosed multiple myeloma subsequently treated with high dose therapy and stem cell transplants termed Total Therapy 2 (pre-treatment TT2) or Total Therapy 3 (pre-treatment TT3).

***Input for Clustering Methods:***

**GE:** scaled gene expression of genes with TPM>1 and standard deviation within the top highest 25% values were used. Total of 2452 genes were used for clustering.

**CNV:** Copy number calls were grouped by cytobands (n=811) using humarray R package. Absolute copy numbers were then categorized as deletion=1 (CN < 2), normal CN=2 (diploid) and gain=3 (CN > 2) for the clustering.

**SNV:** SNV were merged using pathway enrichment analysis using PathScore (48), generating a binary matrix (pathwaysXsamples) for clustering analysis, where 1 was assigned to significant pathways in a sample after if corrected p.value was < 0.05. Number of pathways was 309.

**SV:** Only SVs that were observed in ≥ 5% of the patients were kept for clustering (t(4;14), t(11;14) and t(14;16)).

***Clusters of clusters analysis (COCA):*** ConsensusClusterPlus R-package was used to identify clusters in each data type separately (RNAseq, SNVs, CNVs and LOH) using 1 000 iterations with 70% sample and feature resampling[4]. Number of clusters ranged from 2 to 20 (k=2 to k=20) and hierarchical clustering was used with average innerLinkage and finalLinkage and Pearson correlation as the similarity metric. For the SV data, the number of clusters ranged from 2 to 5 (k=2 to k=5). Clusters were chosen based on cophenetic distances generated from clustering (12 for GE and CNVs, 15 for LOH, 18 for SNVs and 4 for SV data).

COCA's cluster calls from each of the four data types were used to identify relationships between the different classifications. For this purpose, subtypes defined from each platform were coded into a series of indicator variables. Binary matrices were used in ConsensusClusterPlus to identify structure and relationship of the samples. Fourteen clusters were finally identified based on the various datatypes.

***iCluster+:*** This program works with different types of data (categorical (CNVs), continuous (GE) and binary (SNVs and SVs) which allowed a direct use of the distinct datasets. iCluster+ was also run with 70% sample and feature resampling with number of clusters ranging from 2 to 20 clusters (k=2 to k=20). In this case, the number of iterations was 377 the point at which convergence was reached. Lambda parameter was set to 307 (as suggested by the authors) and scaled for the different data types (multinomial=.3, gaussian = .9, binomial=.1). The modal number of clusters selected through the iterations was 12 (40% of the time). A consensus matrix was calculated for the 12 clusters based on sample associations across iterations.

***Selection of clustering algorithm:*** The consensus matrices from the 12 iCluster+ segments and the 14 COCA segments were compared based on similarity within their clusters. Similarity was calculated as the median percentage of times samples grouped together across iterations. iCluster+ showed a significantly higher association within its clusters (t-test p value<.05).

***Identification of master regulators:*** Master regulator (MR) analysis was performed using the msVIPER algorithm in the VIPER R package. Taken as inputs the MM specific regulon model for each candidate regulator protein described above, and the GE profiles associated with MDMS8 and non-MDMS8 samples, the msVIPER algorithm calculated a context-specific protein activity score for each candidate regulator protein by analyzing the enrichment of differential mRNA expression of its regulon genes in MDMS8 versus the rest samples. Each regulator's protein activity score is statistically tested using a null model of random protein activity scores from 1000 permutation of the original GE profiles [5]. Output from the MR analysis was a ranked list of the 20 most activated/inactivated regulator proteins in MDMS8. The top 10 predicted MRs were selected based on FDR<0.01 for further validation.

***Inference of MM-specific gene regulatory network:*** As a prerequisite to the MR analysis, we first applied the hARACNe algorithm [6-8] to reconstruct a gene regulatory network model on a predefined set of candidate regulator proteins using an independent primary MM GE profile (GEP) from GSE2658 [9]. The 6,352 candidate regulator proteins were selected to include all human transcription factors, cofactors and signaling proteins involved in signal transduction via phosphorylation, acetylation, ubiquitination and so on. Before running hARACNe, GEP was first filtered to eliminate all genes with expression interquartile range less than 0.5 and average expression less than 4.5. hARACNe was run up to 3rd order data processing inequality (DPI) and P-value threshold of 1e-8 for mutual information calculation. A consensus network was built from 100 bootstrap runs keeping only edges with a Bonferroni corrected p.value <0.05 based on a Poisson distribution. The output of hARACNe was a gene regulatory network model where each regulator protein was associated with a set of inferred transcriptional targets. We then used the "aracne2regulon" function provided in the VIPER R package in Bioconductor (version 1.16.0) to generate a "regulon" model where each inferred target of a candidate regulator protein was determined to be either "activated" or "repressed", and ambiguous targets were discarded.

***Estimation and exploration of contribution of multi-source omics data to disease segments:*** We utilized the r.jive [10] package 2.1 (https://cran.r-project.org/src/contrib/Archive/r.jive/) to estimate the contribution of each data source in the full dataset of 514 MMRF patients utilized for clustering. Analysis of joint and individual variation (JIVE [10]) showed that >70% of the joint variation of the data was driven by SVs, followed by CNVs (>15%) and GE (>10%) (Supplemental Figure S1C). Interestingly, SNV data, which is commonly used to describe oncogenic drivers in multiple cancers including MM, had the lowest effect on the joint analysis.

### FIGURE LEGENDS

**Supplementary Figure S1. Analytic Methodology.** Left part of the figure shows the process followed in the discovery dataset, with standardized processing of RNAseq (gene expression) and WES data (CNV, SNV and SV), application of two clustering methods, selection of final clusters. Right part of the figure shows validation of HR group in a validation dataset (right), where data was processed using the same methodology than in the discovery dataset.

**Supplementary Figure S2. Consensus Matrix and Joint Variation Analysis. A)** Consensus matrix (samples by samples matrix sorted equally in rows and columns), representing how many times every two samples were clustered together across the iterations. Color scheme goes from yellow (100% of the iterations the two samples were grouped together) to red (never the two samples were grouped together). **B)** Jive figure showing the joint, individual and residual variation across data types. **C)** Figure showing the data variation explained per number of clusters.

**Supplementary Figure S3. MAF expression. A)** MAF normalized gene expression in patients without (0) and with (1) t(14;16). **B)** MAF normalized gene expression across the 12 clusters.

**Supplementary Figure S4. Genomic markers in MDMS8 versus non-MDMS8 samples. A)** PFS (left) and OS (right) Kaplan Meier figures for MDMS8 gain1q vs non-MDMS8 gain1q patients (top). Table listing up-regulated gene expression pathways of gain1q in MDMS8 versus rest of gain1q patients (bottom); **B)** PFS (left) and OS (right) Kaplan Meier figures for MDMS8 t(4;14) vs non-MDMS8 t(4;14) patients (top). Table listing up-regulated gene expression pathways of t(4;14) in MDMS8 versus rest of gain1q patients (bottom); **C)** PFS (left) and OS (right) Kaplan Meier figures for MDMS8 t(14;16) vs non-MDMS8 t(14;16) patients; **D)** PFS (left) and OS (right) Kaplan Meier figures for MDMS8 del13q vs non-MDMS8 del13q patients; **E)** PFS (left) and OS (right) Kaplan Meier figures for MDMS8 del17p vs non-MDMS8 del17p patients.

**Supplementary Figure S5. DNA damage pathway scores in MDMS8 vs rest.** Boxplot figures for the Base Excision Repair (BER), Nucleotide Excision Repair (NER) , Mistmatch Repair (MMR), the Fanconi anemia (FA), non-homologous end joining (NHEJ), Nucleotide Pool Maintenance (NP), Direct Damage reversal/repair (DR) and translesion DNA repair (TLS).

**Supplementary Figure S6. Validation of MDMS8 in an Independent Dataset (IFM). A)** Kaplan-Meier curves for OS of MDMS-8 like (patients selected by the GE linear model classifier) vs non-MDMS8-like patients in the IFM (left) and APEX (right) validation cohorts; **B)** Validation of increase of genomic loss in MDMS8-like patients in IFM dataset.

**Supplementary Figure S7. Comparison of MDMSs to Segments from Broyl et al.** Gene expression enrichment of Broyl et al GE patient subgroups signatures[11] across the twelve molecularly defined myeloma segments. Red represents positive enrichment; blue represents negative enrichment. Blue squares highlight significant association (enrichment scores t-test p<0.01) between the Broyl et al signatures and MDMS disease segments.

**Supplementary Figure S8. Comparison to other high-risk GE signatures.** Venn diagram showing the patient populations selected on the discovery dataset by the multiple myeloma high-risk gene expression signatures UAMS70 and EMC92+ISS III and MDMS8.

**Supplementary Figure S9. Master regulators of high-risk MDMS8.** Left side of the plot shows the distribution of the positively (red) and negatively (blue) correlated targets of the MR on the gene list ranked by differential mRNA expression in MDMS8 compared with the rest. The right side of the plot shows fold change of inferred activity and mRNA expression level of the MRs in MDMS8 vs rest of the samples.

**Supplementary Figure S1**

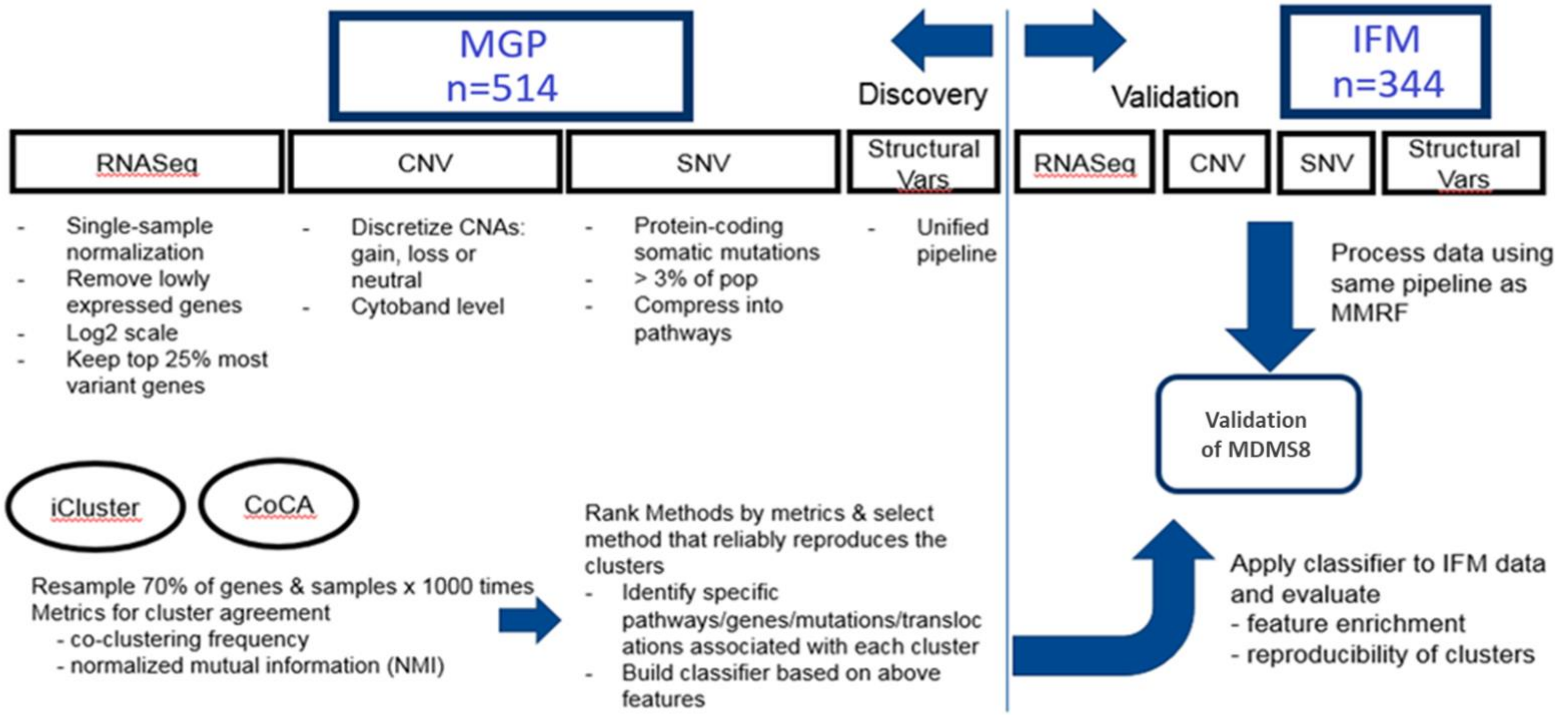

**Supplementary Figure S2**

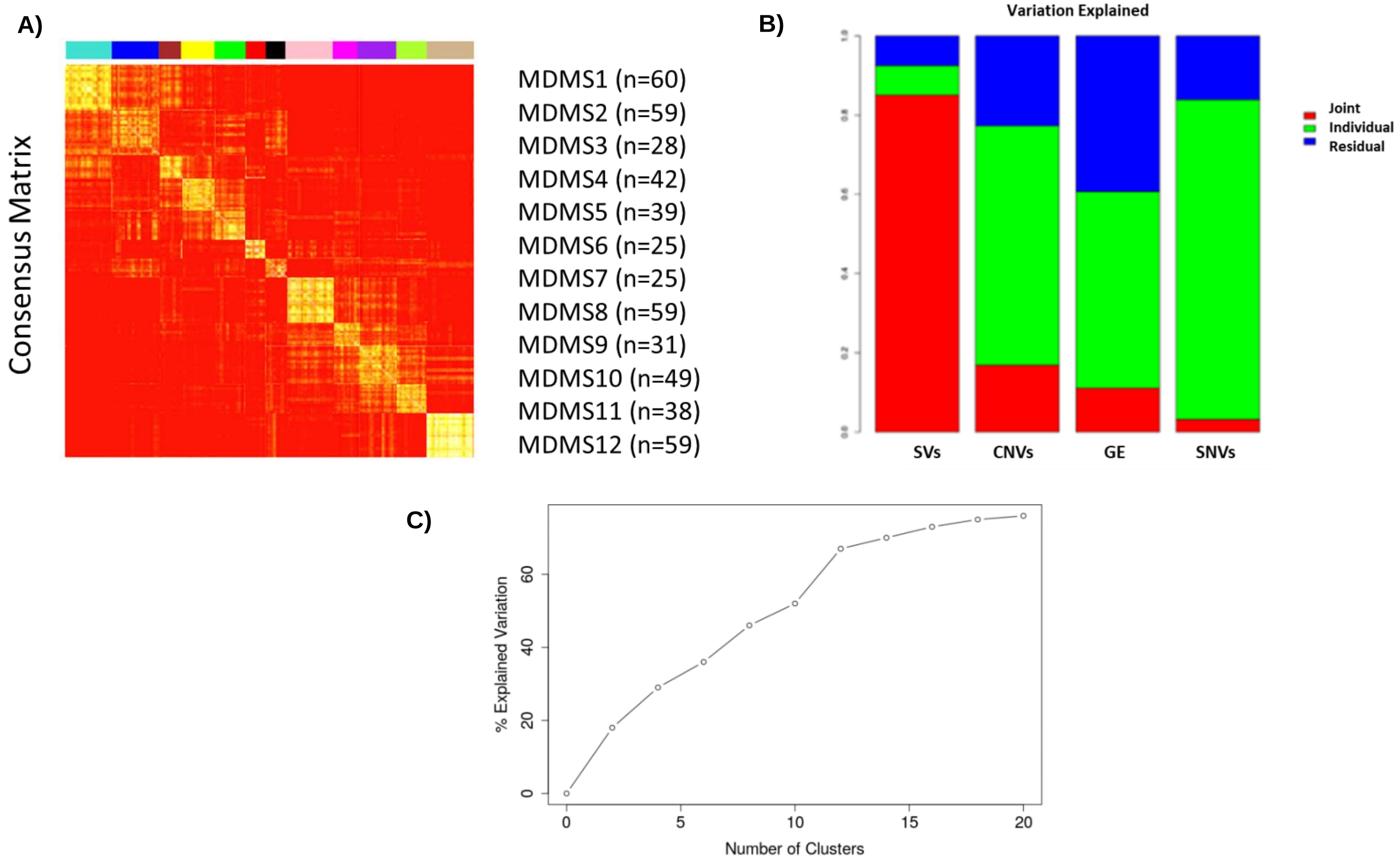

**Supplementary Figure S3**

**A)**

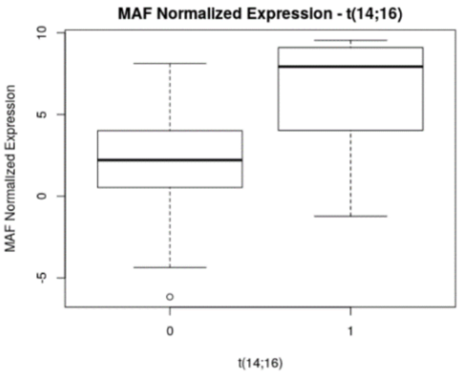


**B)**

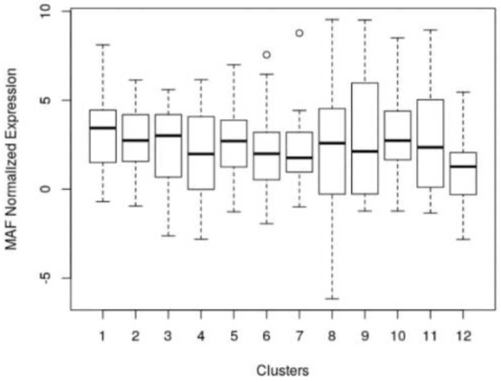

**Supplementary Figure S4-A**

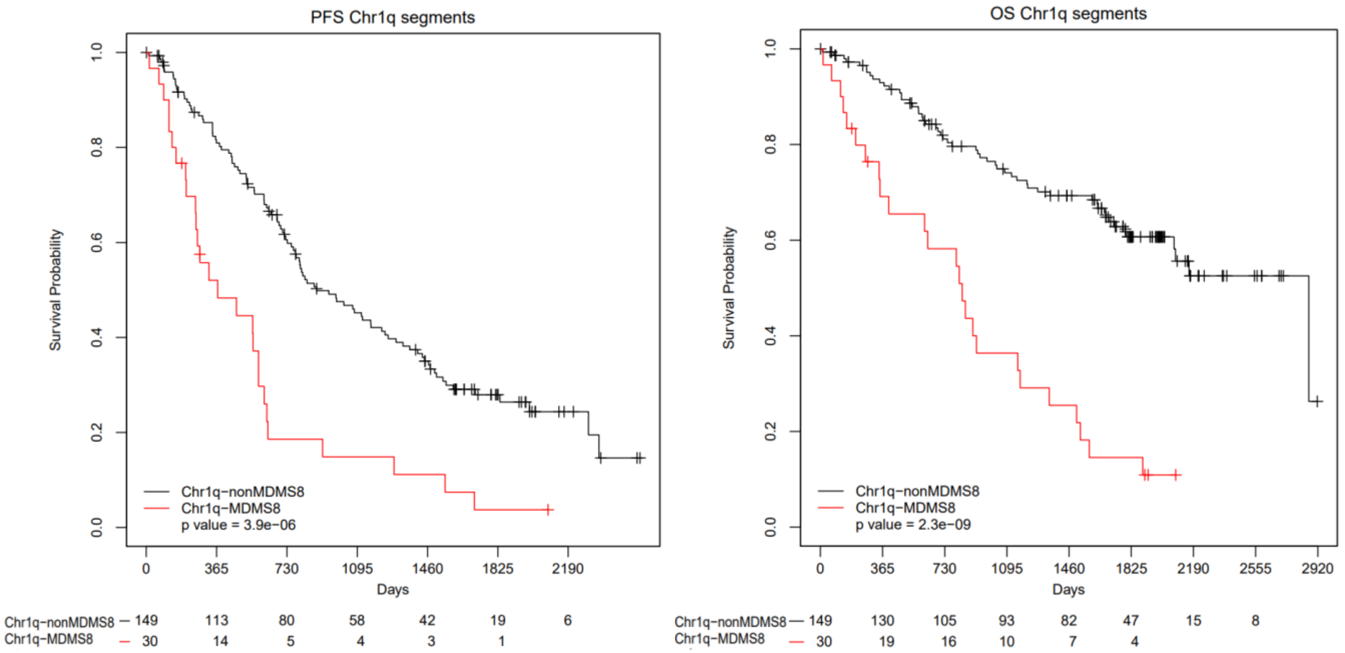


| ID | NES | p.adjust |
|---|---|---|
| HALLMARK_MYC_TARGETS_V2 | 2.509755 | 0.0152532 |
| HALLMARK_PI3K_AKT_MTOR_SIGNALING | 1.923333 | 0.0152532 |
| HALLMARK_UV_RESPONSE_UP | 1.766947 | 0.0152532 |
| HALLMARK_DNA_REPAIR | 1.754434 | 0.0152532 |
| HALLMARK_OXIDATIVE_PHOSPHORYLATION | 1.859213 | 0.0152532 |
| HALLMARK_APOPTOSIS | 2.026959 | 0.0152532 |
| HALLMARK_TNFA_SIGNALING_VIA_NFKB | 1.808707 | 0.0152532 |
| HALLMARK_G2M_CHECKPOINT | 2.524248 | 0.0152532 |
| HALLMARK_MTORC1_SIGNALING | 2.152247 | 0.0152532 |
| HALLMARK_E2F_TARGETS | 2.969838 | 0.0152532 |
| HALLMARK_MYC_TARGETS_V1 | 3.076199 | 0.0152532 |
| HALLMARK_P53_PATHWAY | 1.597128 | 0.0263713 |

**Supplementary Figure S4-B**

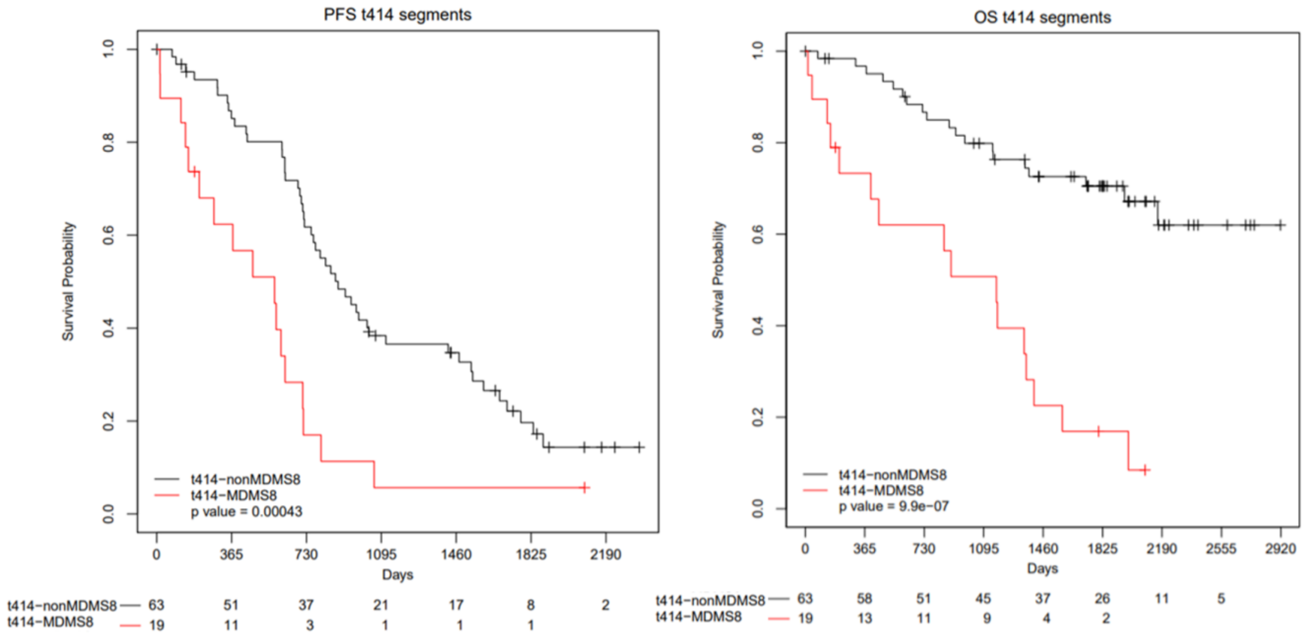


| ID | NES | p.adjust |
|---|---|---|
| HALLMARK_MYC_TARGETS_V2 | 2.361483 | 0.016469 |
| HALLMARK_E2F_TARGETS | 3.451184 | 0.016469 |
| HALLMARK_MITOTIC_SPINDLE | 2.281763 | 0.016469 |
| HALLMARK_MYC_TARGETS_V1 | 2.239475 | 0.016469 |
| HALLMARK_SPERMATOGENESIS | 2.003444 | 0.016469 |
| HALLMARK_G2M_CHECKPOINT | 3.288724 | 0.016469 |

**Supplementary Figure S4-C**

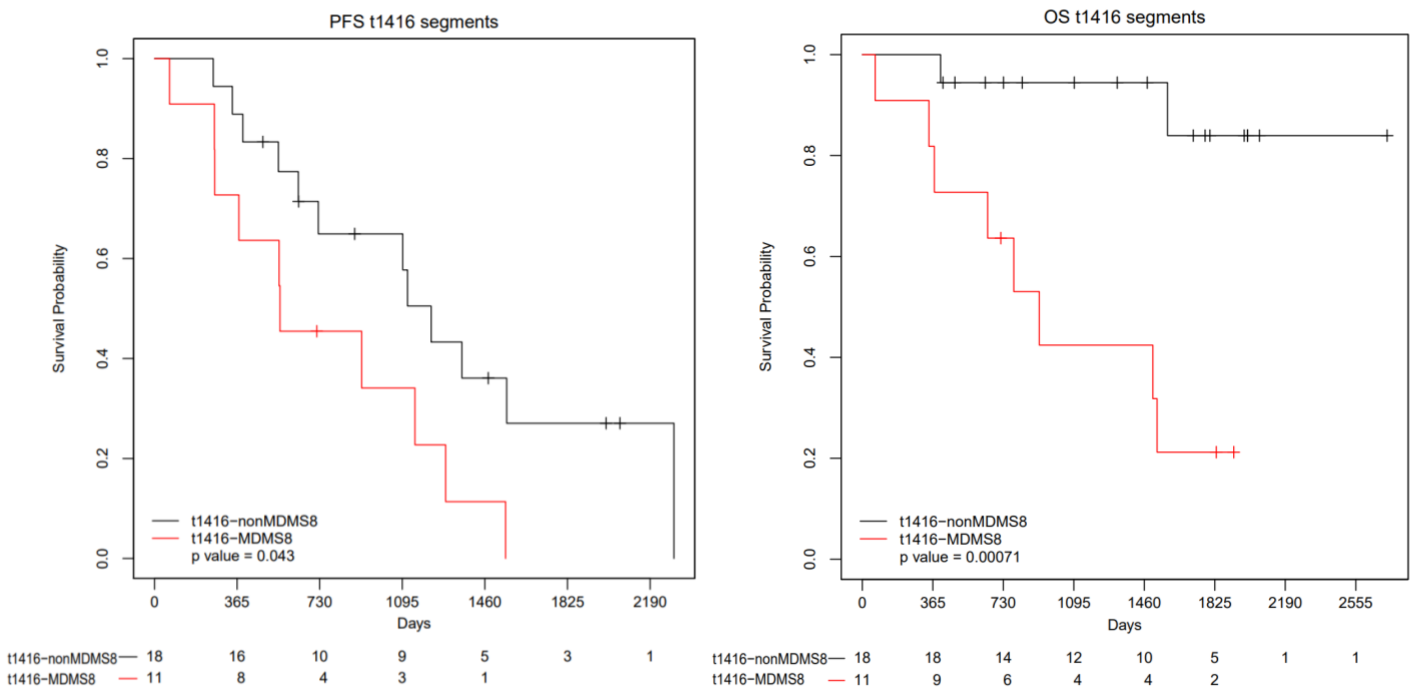

**Supplementary Figure S4-D**

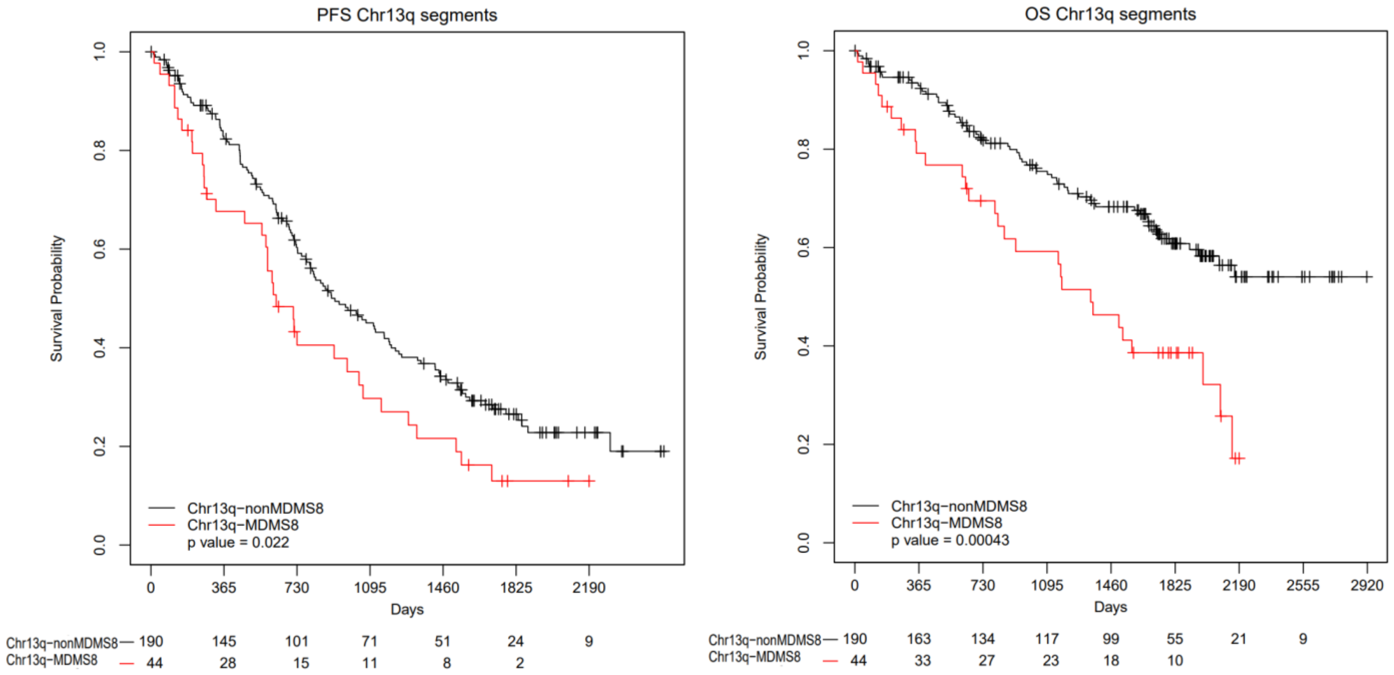

**Supplementary Figure S4-E**

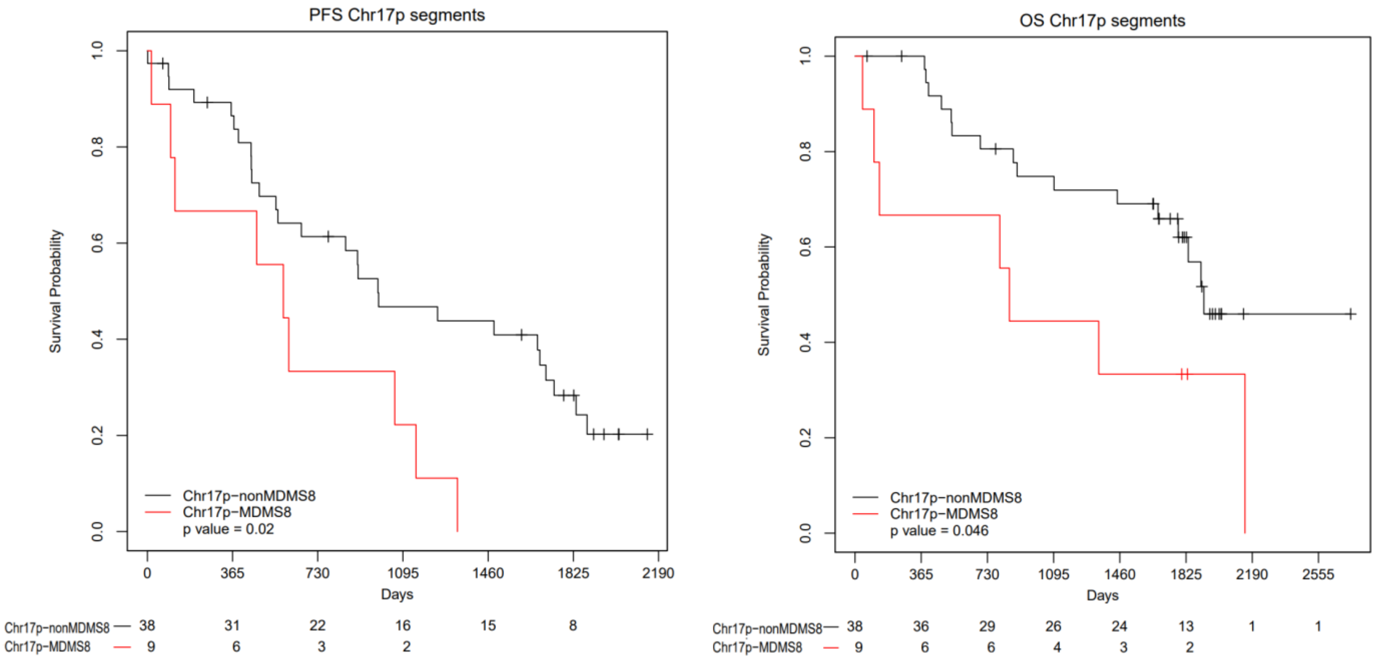

### Supplementary Figure S5

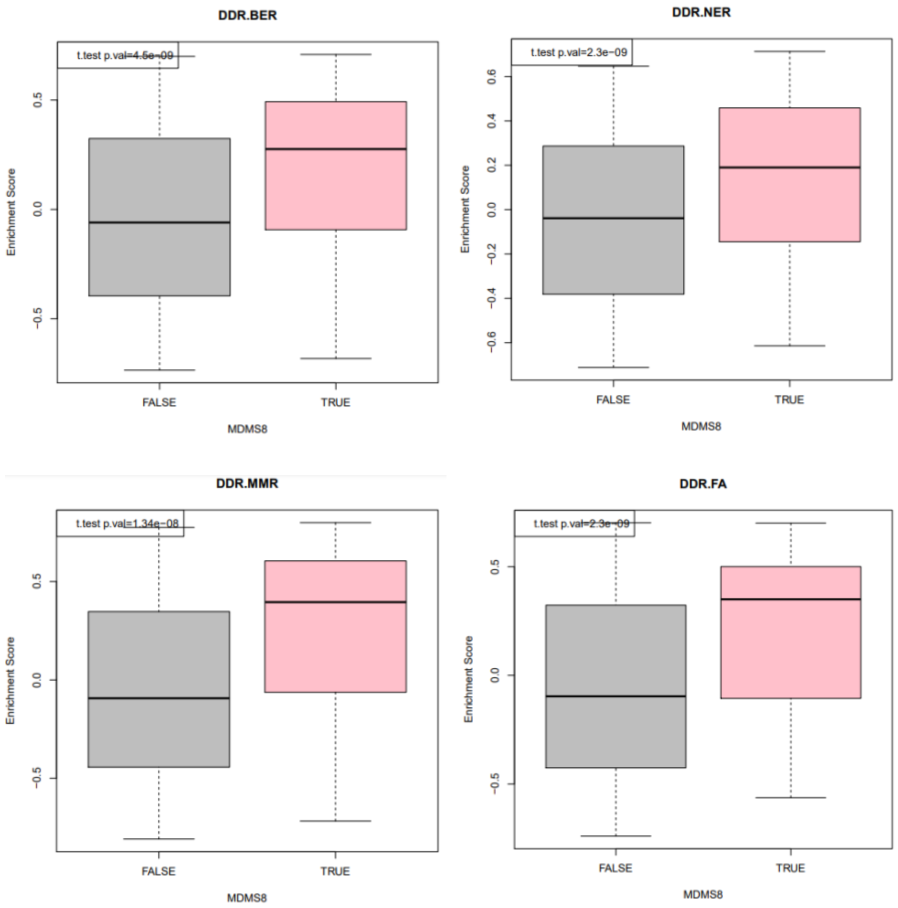

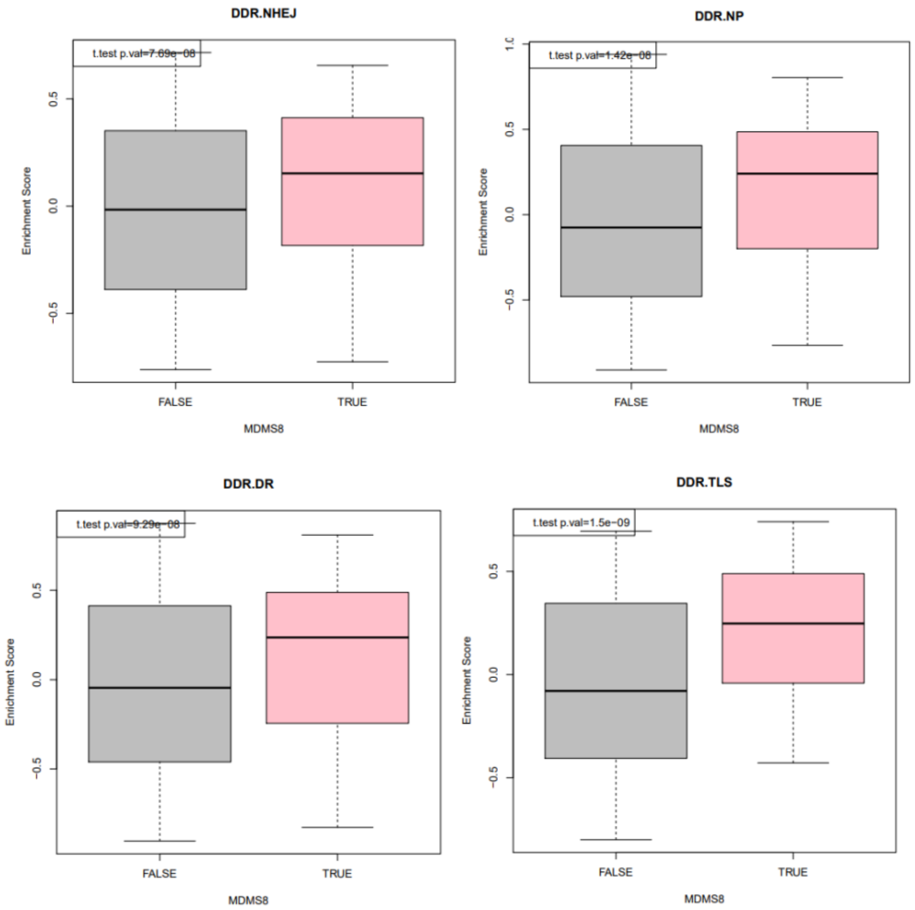
DDR.NHEJ
t.test p.val=7.69e−08
Enrichment Score
MDMS8
FALSE
TRUE
DDR.NP
t.test p.val=1.42e−08
Enrichment Score
MDMS8
FALSE
TRUE
DDR.DR
t.test p.val=9.29e−08
Enrichment Score
MDMS8
FALSE
TRUE
DDR.TLS
t.test p.val=1.5e−09
Enrichment Score
MDMS8
FALSE
TRUE

### Supplementary Figure S6-A

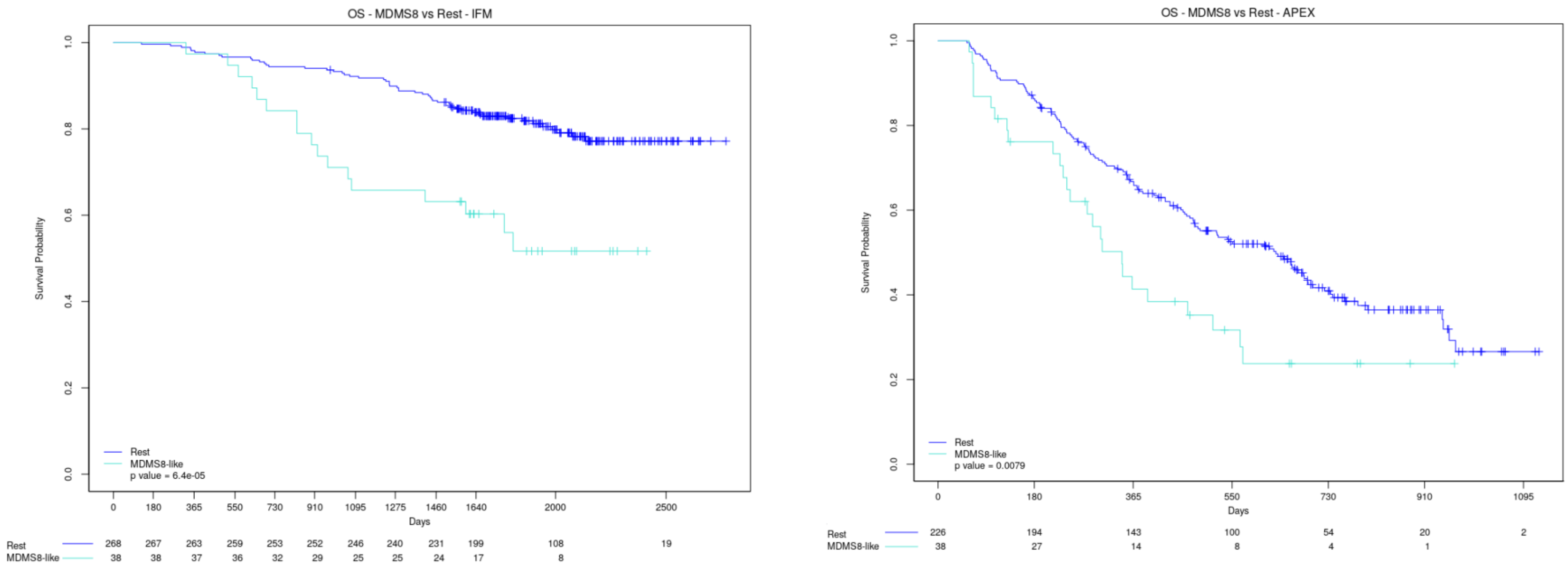

**Supplementary Figure S6-B**

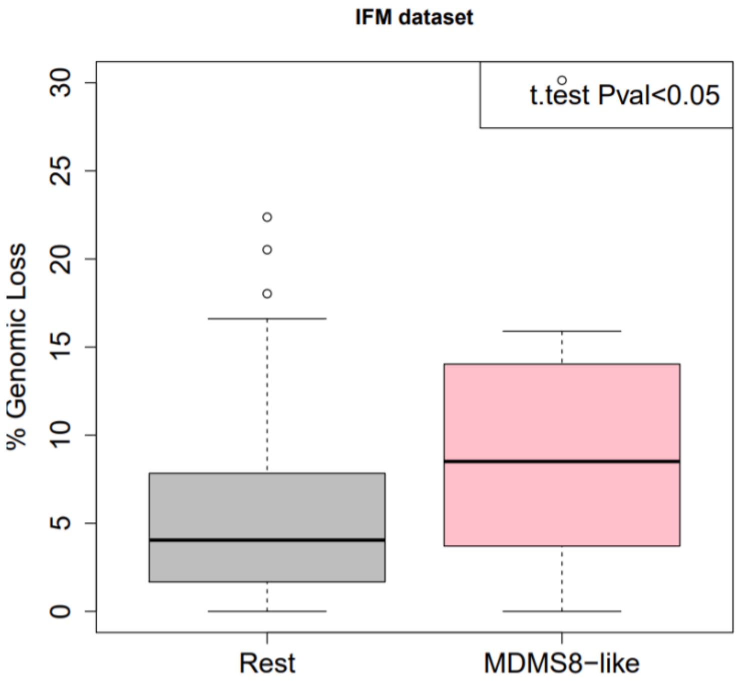

**Supplementary Figure S7**

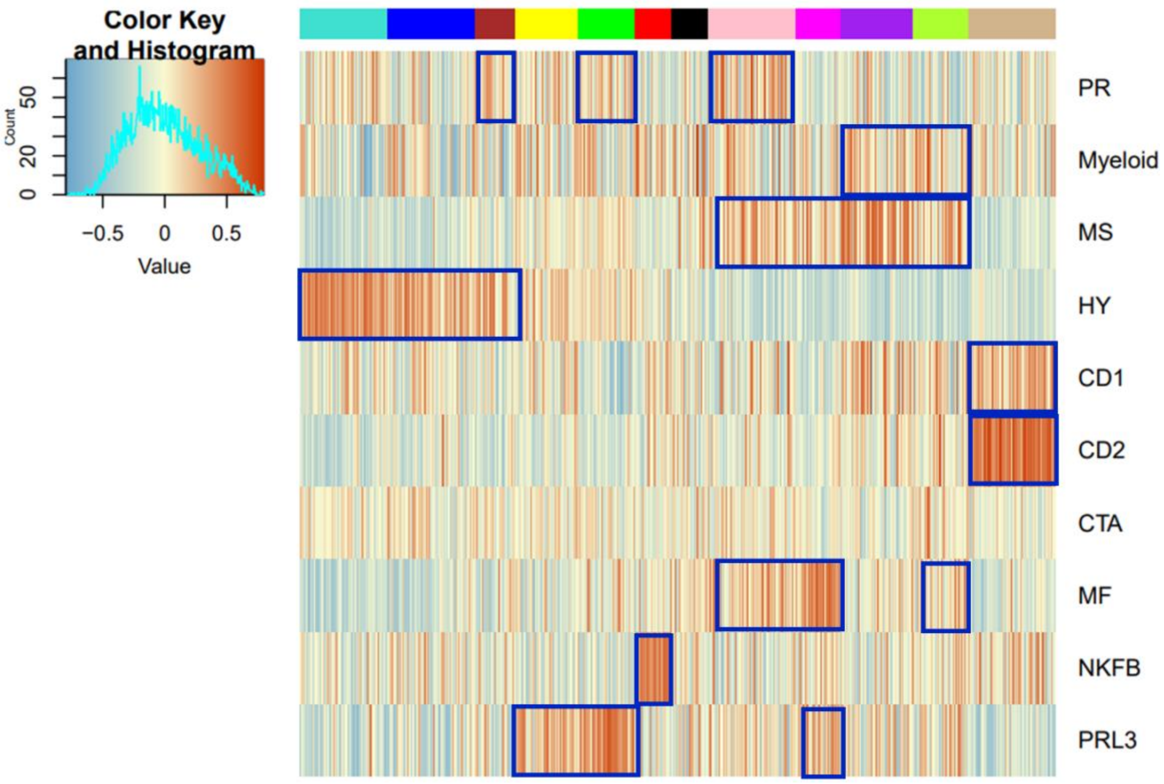

**Supplementary Figure S8**

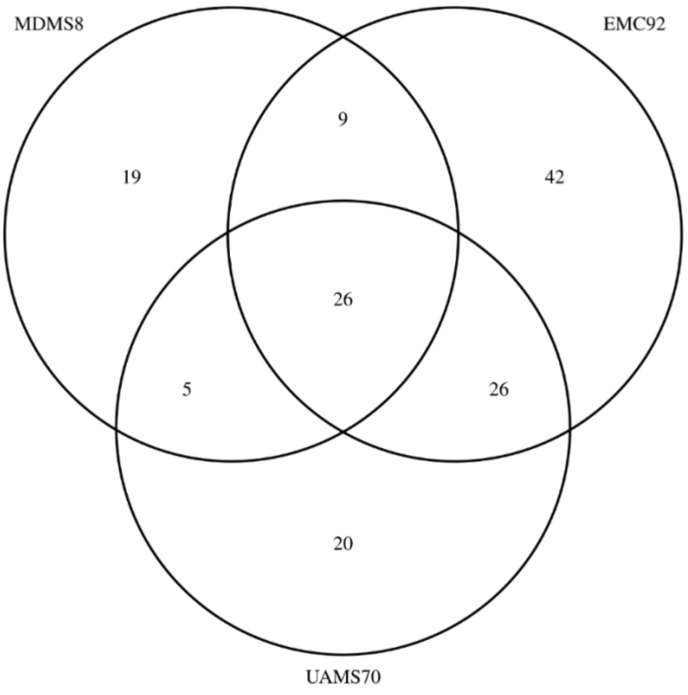

**Supplementary Figure S9**

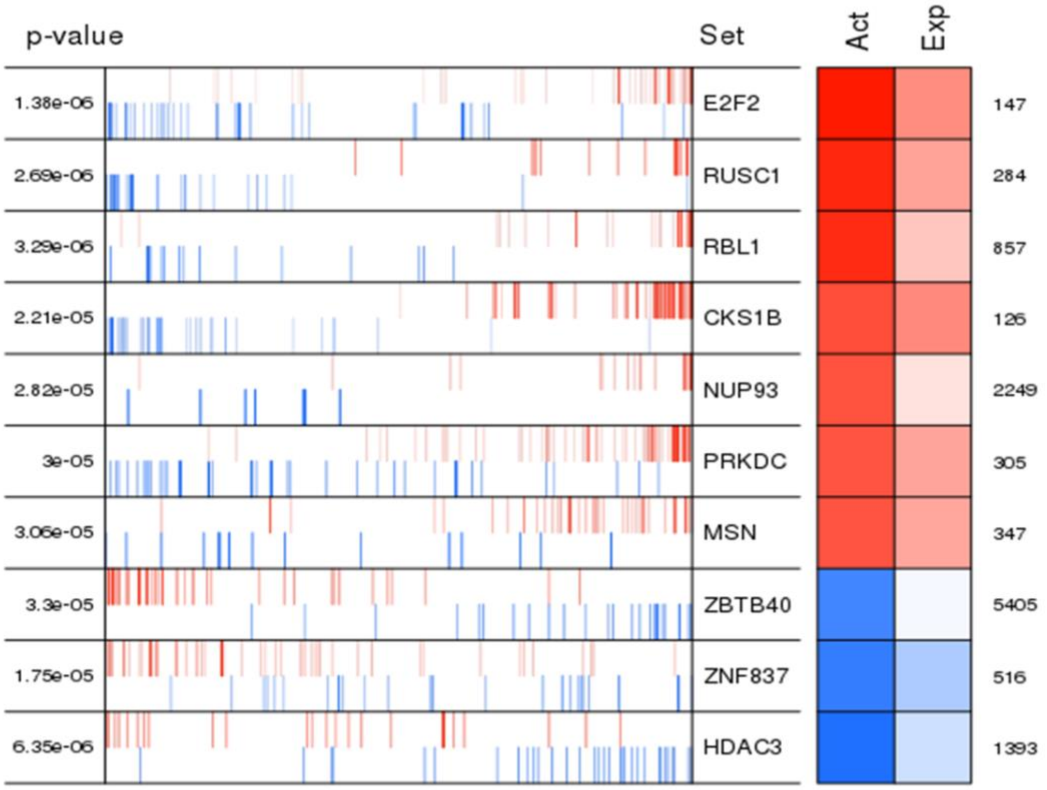


### TABLE LEGENDS

**Supplementary Table S1. Dataset Characteristics.** This table contains number of patients, median OS, PFS and age and prevalence of ISS III in the three datasets used in this work (subset of MMRF, IFM and APEX).

**Supplementary Table S2. COCA vs iCluster+ segments.** Table showing consensus between COCA clusters (columns) and iCluster+ (rows). Highlighted in red the highest overlaps between the results.

**Supplementary Table S3. Cohort and Genomic Features of MDMS1–12.** Table listing clinical, genomic and transcriptomic characteristics of the molecularly defined segments. Features in this table included previously known markers of MM. Nominal p.values were calculated by t.test for continuous values and fisher exact test for binary values. Highlighted in red significant nominal p.values ($p < .05$).

**Supplementary Table S4. Features Summary.** This table contains all the features used for the clustering work and their prevalence (for single nucleotide variants, copy number variants and structural variants) and their medians (for gene expression).

**Supplementary Table S5. Confusion matrix of cluster labels and classifier predictions on the discovery dataset.** This table contains all the features used for the clustering work and their prevalence (for single nucleotide variants, copy number variants and structural variants) and their medians (for gene expression).

**Supplementary Table S6. Comparison of MDMS8 to the previously defined GE UAMS groups.** This table contains a comparison of our MDMS1-MDMS12all the features used for the clustering work and their prevalence (for single nucleotide variants, copy number variants and structural variants) and their medians (for gene expression).

**Supplementary Table S1. Characteristics of Discovery and Validation Cohorts**

| | Discovery (MMRF IA17 subset) | MMRF IA17 | Validation of MDMS8 (IFM 2009) |
|---|---|---|---|
| N | 514 | 970 | 344 |
| Median PFS (95% CI, mo) | 34.1 (28.6, 38.0) | 32.9 (30.0, 36.6) | 43.4 (40.3, NE) |
| Median OS (95% CI, mo) | 94.0 (71.5, NR) | NE (64.3, NE) | NE (NE, NE) |
| Median Age (yr) | 65 | 63 | 59 |
| ISS3 (%) | 29.3% | 26.2% | 20.9% |

**Supplementary Table S1. COCA and iCluster+ Segmentation**

| | | COCA | | | | | | | | | | | | | |
|---|---|---|---|---|---|---|---|---|---|---|---|---|---|---|---|
| | | **1** | **2** | **3** | **4** | **5** | **6** | **7** | **8** | **9** | **10** | **11** | **12** | **13** | **14** |
| **iCluster+** | **1** | 15 | 5 | 0 | 0 | 1 | 0 | 7 | 0 | 9 | 0 | 0 | 0 | 12 | 0 |
| | **2** | 0 | 5 | 0 | 2 | 6 | 2 | 3 | 0 | 4 | 0 | 0 | 0 | 2 | 1 |
| | **3** | 0 | 3 | 23 | 1 | 1 | 0 | 0 | 20 | 1 | 9 | 0 | 0 | 0 | 2 |
| | **4** | 0 | 0 | 1 | 18 | 0 | 14 | 0 | 4 | 0 | 0 | 2 | 3 | 0 | 0 |
| | **5** | 0 | 4 | 3 | 6 | 3 | 1 | 1 | 21 | 2 | 6 | 0 | 10 | 2 | 0 |
| | **6** | 0 | 1 | 0 | 16 | 2 | 14 | 1 | 0 | 1 | 1 | 1 | 2 | 0 | 0 |
| | **7** | 2 | 4 | 0 | 0 | 1 | 4 | 10 | 0 | 14 | 0 | 0 | 1 | 1 | 1 |
| | **8** | 4 | 2 | 0 | 0 | 1 | 0 | 17 | 0 | 17 | 0 | 12 | 0 | 6 | 0 |
| | **9** | 1 | 0 | 0 | 0 | 0 | 0 | 6 | 0 | 49 | 0 | 0 | 1 | 2 | 0 |
| | **10** | 0 | 2 | 5 | 2 | 3 | 0 | 0 | 4 | 0 | 2 | 4 | 3 | 0 | 3 |
| | **11** | 0 | 4 | 5 | 2 | 0 | 0 | 3 | 1 | 1 | 0 | 0 | 8 | 0 | 1 |
| | **12** | 13 | 3 | 0 | 1 | 0 | 1 | 4 | 0 | 3 | 0 | 5 | 0 | 1 | 0 |

### Supplementary Table S2. Cohort and Genomic Characteristics of MDMS 1-12 (p-values)

| | | MDMS1 (n=60) | MDMS2 (n=59) | MDMS3 (n=28) | MDMS4 (n=42) | MDMS5 (n=39) | MDMS6 (n=25) | MDMS7 (n=25) | MDMS8 (n=59) | MDMS9 (n=31) | MDMS10 (n=49) | MDMS11 (n=38) | MDMS12 (n=59) |
|---|---|---|---|---|---|---|---|---|---|---|---|---|---|
| Cohort Characteristics | ISS stage III | 0.772 | 0.957 | 0.062 | 0.086 | 0.477 | 0.632 | 0.53 | <0.001 | 0.967 | 0.842 | 0.603 | 0.999 |
| | median PFS | 0.054 | 0.6 | 0.46 | 0.353 | 0.35 | 0.98 | 0.1 | <1e-5 | 0.08 | 0.812 | 0.88 | 0.3 |
| | median OS | 0.45 | 0.47 | 0.11 | 0.13 | 0.95 | 0.7 | 0.18 | <1e-8 | 0.2 | 0.95 | 0.15 | 0.48 |
| | biallelic TP53 | 0.576 | 0.862 | 0.215 | 1 | 1 | 0.18 | 1 | 0.027 | 0.25 | 1 | 0.713 | 0.862 |
| Structural Variants | t(4;14) | 1 | 0.998 | 0.993 | 0.977 | 0.999 | 1 | 0.586 | <0.001 | 0.102 | <0.001 | 0.026 | 1 |
| | t(11;14) | 0.999 | 0.81 | 0.996 | 1 | 1 | 0.763 | 0.235 | 0.996 | 1 | 0.535 | 0.097 | <0.001 |
| | t(14;16) | 0.975 | 0.974 | 0.812 | 0.922 | 1 | 0.774 | 0.774 | <0.001 | <0.001 | 0.95 | 0.899 | 1 |
| Signaling Pathways | CELL_CYCLE | 0.251 | <0.001 | 0.005 | 0.054 | 0.909 | <0.001 | 0.321 | <0.001 | 0.184 | 0.392 | 0.431 | 0.204 |
| | CC_CHECKPOINTS | 0.741 | 0.003 | 0.005 | 0.765 | 0.755 | <0.001 | 0.208 | <0.001 | 0.783 | 0.283 | 0.161 | 0.132 |
| | DNA_REPAIR | 0.086 | 0.168 | 0.008 | 0.635 | 0.833 | <0.001 | 0.285 | <0.001 | 0.981 | 0.018 | 0.143 | 0.227 |
| | INTERFERON | <0.001 | 0.751 | <0.001 | 0.013 | 0.409 | 0.161 | 0.075 | 0.826 | 0.357 | 0.032 | 0.01 | 0.123 |
| Copy Number Deletions | del1p22.1 | 0.792 | 0.863 | 0.022 | 0.97 | 0.951 | 0.282 | 0.463 | <0.001 | 0.347 | 0.501 | 0.602 | 1 |
| | del13q14.3 | 1 | 1 | 0.809 | 0.999 | 0.005 | 0.947 | 0.639 | <0.001 | <0.001 | <0.001 | 0.341 | 1 |
| | del14q32.33 | 0.323 | 0.215 | 0.845 | 0.286 | 0.361 | 0.193 | 0.612 | 0.345 | 0.025 | 0.278 | 0.731 | 1 |
| | del16q24.1 | 0.407 | 0.744 | 0.008 | 0.931 | 0.793 | 0.136 | 0.062 | 0.003 | 0.023 | 1 | 0.873 | 0.999 |
| | del17p13.1 | 0.666 | 0.074 | 0.101 | 0.916 | 0.716 | 0.405 | 0.687 | 0.074 | 0.312 | 1 | 0.698 | 0.981 |
| Copy Number Gains (CNV ≥ 3) | gain 1q21.3 | 1 | 1 | 0.533 | 0.511 | 0.001 | 0.97 | 0.693 | 0.005 | <0.001 | 0.082 | 0.727 | 0.963 |
| | gain 3q26.2 | 0.108 | <0.001 | 0.018 | <0.001 | 0.014 | 0.066 | 0.391 | 1 | 0.999 | 1 | 1 | 1 |
| | gain 5q33.2 | <0.001 | <0.001 | <0.001 | <0.001 | <0.001 | 0.123 | 1 | 1 | 1 | 1 | 1 | 1 |
| | gain 11q23.3 | <0.001 | <0.001 | <0.001 | 0.951 | 1 | 0.012 | 0.272 | 1 | 1 | 1 | 0.999 | 0.899 |
| | gain 15q25.1 | <0.001 | <0.001 | <0.001 | <0.001 | <0.001 | 0.045 | 0.993 | 1 | 1 | 1 | 1 | 1 |
| Driver Mutations | *ARID2* | 0.979 | 0.881 | 0.823 | 0.928 | 0.02 | 0.786 | 1 | 0.004 | 1 | 0.561 | 0.385 | 0.693 |
| | *ATM* | 0.499 | 0.83 | 0.759 | 0.922 | 0.505 | 0.699 | 0.919 | 0.305 | 0.005 | 0.861 | 0.275 | 0.83 |
| | *BIRC2* | 1 | 0.708 | 1 | 1 | 1 | 1 | 1 | 0.019 | 0.117 | 0.636 | 0.539 | 0.708 |
| | *CCND1* | 0.989 | 0.988 | 0.869 | 0.955 | 0.507 | 0.52 | 1 | 0.201 | 0.895 | 0.232 | 0.938 | <0.001 |
| | *DIS3* | 0.97 | 0.998 | 0.902 | 0.933 | 0.399 | 1 | 0.216 | 0.515 | 0.003 | 0.006 | 0.898 | 0.126 |
| | *EGR1* | 0.767 | 0.984 | 0.852 | 0.945 | 0.008 | 0.211 | 0.817 | 0.539 | 0.32 | 0.629 | 0.722 | 0.539 |
| | *FAM46C* | <0.001 | 0.339 | 0.054 | 0.817 | 0.973 | 0.646 | 1 | 0.904 | 0.582 | 0.992 | 0.969 | 0.496 |
| | *FGFR3* | 0.918 | 0.984 | 1 | 0.945 | 0.932 | 0.817 | 0.817 | 0.325 | 0.32 | <0.001 | 0.027 | 0.984 |
| | *IRF4* | 0.964 | 0.961 | 0.162 | 1 | 0.878 | 1 | 0.735 | 0.348 | 0.201 | 0.46 | 0.589 | 0.021 |
| | *MAX* | 0.959 | 0.956 | 1 | 0.624 | 0.582 | 0.347 | 1 | 0.017 | 0.797 | 0.433 | 0.03 | 0.956 |
| | *NF1* | 0.646 | 0.802 | 0.737 | 0.984 | 0.048 | 0.391 | 0.062 | 0.979 | 0.95 | 0.452 | 0.245 | 0.437 |
| | *NRAS* | 0.033 | 0.315 | 0.848 | 0.105 | 0.419 | 0.357 | 1 | 0.607 | 1 | 1 | 0.88 | 0.012 |
| | *PRKD2* | 0.968 | 0.84 | 1 | 0.906 | 0.337 | 0.749 | 0.383 | 0.376 | 0.822 | 0.009 | 0.61 | 0.619 |
| | *RB1* | 1 | 0.855 | 0.058 | 0.687 | 0.646 | 1 | 1 | 0.031 | 0.234 | 0.769 | 0.343 | 0.645 |
| | *SETD2* | 0.814 | 0.956 | 1 | 0.888 | 0.582 | 0.347 | 1 | 0.017 | 0.797 | 0.078 | 0.567 | 0.806 |
| | *TGDS* | 1 | 0.366 | 1 | 0.612 | 0.044 | 1 | 1 | 1 | 1 | 0.078 | 0.574 | 0.742 |
| | *TRAF3* | 0.959 | 0.741 | 0.554 | 0.943 | 0.78 | 1 | 0.097 | 0.025 | 0.028 | 0.374 | 0.337 | 0.989 |

### Supplementary Table S4. Classifier features

| Features | Coefficients | Features | Coefficients |
|---|---|---|---|
| (Intercept) | 0.009053312 | CCL19 | 0.155221827 |
| CCL5 | 0.385757202 | FCGR2C | -0.101665317 |
| RRBP1 | -0.078819425 | TRHR | -0.236674164 |
| DKK3 | 0.031071851 | TRAV12-2 | 0.016426865 |
| CLDN3 | -0.004582616 | MBOAT2 | 0.008170334 |
| IRS1 | -0.384203111 | SFTPD | -0.291655315 |
| SGCE | 0.031558992 | NAMPT | 0.037437401 |
| MLF1 | 0.158155078 | KLF11 | -0.081413100 |
| HLA-DOB | 0.182813660 | SIGIRR | 0.018719064 |
| BCL2A1 | 0.045511486 | DNMT3L | -0.124415489 |
| GABRR1 | -0.135534412 | LRRTM4 | -0.023328024 |
| TULP1 | -0.011410561 | LMAN1L | -0.122773804 |
| TNFRSF11A | 0.293801060 | CABP5 | 0.116992702 |
| LHCGR | 0.011974565 | AFAP1-AS1 | -0.148885221 |
| ID3 | 0.012311308 | CACHD1 | -0.105518650 |
| IL13 | -0.073307312 | EBF1 | 0.007314996 |
| KRT9 | 0.098836702 | LINC00324 | 0.295082102 |
| GRAP2 | 0.268433270 | TPPP2 | -0.075230309 |
| LGALS2 | 0.010319057 | RBM24 | 0.109861626 |
| GRIA1 | -0.120511871 | C6orf52 | 0.017618441 |
| GPR26 | -0.159368733 | RBP7 | 0.030778465 |

### Supplementary Table S5. Confusion matrix of cluster labels and classifier predictions on the discovery dataset

| | | | | | | | Cluster | Labels | | | | | | | |
|---|---|---|---|---|---|---|---|---|---|---|---|---|---|---|---|
| | | **MDMS1** | **MDMS2** | **MDMS3** | **MDMS4** | **MDMS5** | **MDMS6** | **MDMS7** | **MDMS8** | **MDMS9** | **MDMS10** | **MDMS11** | **MDMS12** | **Precision** | |
| | **MDMS1** | 170 | 63 | 27 | 8 | 0 | 3 | 1 | 0 | 0 | 5 | 0 | 0 | 0,61 | |
| **P** | **MDMS2** | 42 | 121 | 13 | 10 | 3 | 9 | 10 | 1 | 0 | 0 | 0 | 0 | 0,58 | |
| **R** | **MDMS3** | 6 | 10 | 23 | 6 | 8 | 0 | 0 | 3 | 1 | 0 | 0 | 0 | 0,40 | |
| **E** | **MDMS4** | 6 | 7 | 3 | 92 | 27 | 1 | 8 | 2 | 5 | 0 | 3 | 0 | 0,60 | |
| **D** | **MDMS5** | 3 | 22 | 12 | 31 | 88 | 3 | 9 | 2 | 5 | 0 | 10 | 0 | 0,48 | |
| **I** | **MDMS6** | 0 | 3 | 0 | 1 | 0 | 69 | 0 | 1 | 0 | 0 | 1 | 6 | 0,85 | |
| **C** | **MDMS7** | 0 | 2 | 2 | 0 | 6 | 0 | 17 | 4 | 4 | 0 | 5 | 0 | 0,43 | |
| **T** | **MDMS8** | 0 | 2 | 6 | 3 | 1 | 9 | 1 | 176 | 9 | 12 | 5 | 5 | 0,77 | |
| **I** | **MDMS9** | 0 | 0 | 9 | 0 | 16 | 0 | 13 | 5 | 86 | 0 | 13 | 11 | 0,56 | |
| **O** | **MDMS10** | 0 | 3 | 0 | 0 | 4 | 1 | 11 | 15 | 8 | 150 | 30 | 7 | 0,66 | |
| **N** | **MDMS11** | 0 | 1 | 0 | 4 | 1 | 0 | 8 | 5 | 6 | 20 | 71 | 3 | 0,60 | |
| | **MDMS12** | 0 | 8 | 0 | 4 | 0 | 0 | 10 | 3 | 0 | 23 | 15 | 204 | 0,76 | **Median Recall** |
| | | 0,75 | 0,50 | 0,24 | 0,58 | 0,57 | 0,73 | 0,19 | 0,81 | 0,69 | 0,71 | 0,46 | 0,86 | **Median Precision** | 0,64 |
| | | | | | | | **Recall** | | | | | | | 0,60 | |

**Supplementary Table S6. Comparison of MDMS8 to the previously defined GE UAMS groups**

| | CD1 | CD2 | HY | LB | MF | MS | MY | PR |
|---|---|---|---|---|---|---|---|---|
| **MDMS1** | 0 | 1 | 35 | 1 | 0 | 0 | 11 | 0 |
| **MDMS2** | 2 | 5 | 51 | 3 | 0 | 0 | 42 | 1 |
| **MDMS3** | 0 | 0 | 8 | 1 | 0 | 1 | 5 | 4 |
| **MDMS4** | 1 | 1 | 3 | 7 | 0 | 0 | 13 | 13 |
| **MDMS5** | 0 | 2 | 2 | 12 | 1 | 3 | 16 | 3 |
| **MDMS6** | 1 | 0 | 0 | 0 | 1 | 0 | 3 | 0 |
| **MDMS7** | 1 | 2 | 0 | 7 | 0 | 1 | 2 | 0 |
| **MDMS8** | 3 | 2 | 1 | 4 | 11 | 19 | 13 | 20 |
| **MDMS9** | 0 | 0 | 0 | 8 | 13 | 6 | 6 | 0 |
| **MDMS10** | 8 | 5 | 0 | 0 | 0 | 21 | 6 | 0 |
| **MDMS11** | 3 | 3 | 0 | 4 | 6 | 9 | 14 | 0 |
| **MDMS12** | 6 | 36 | 2 | 1 | 0 | 0 | 4 | 0 |
